# DNA damage-induced YTHDC1 O-GlcNAcylation promotes homologous recombination by enhancing N6-methyladenosine binding

**DOI:** 10.1101/2022.09.03.506498

**Authors:** Mengyao Li, Jie Li, Aiyun Yuan, Weidong Dong, Suwei Dong, Yun-Gui Yang, Yibo Wang, Chen Wu, Jing Li

**Affiliations:** School of Life Sciences, Institute of Life Sciences and Green Development, Hebei University, Baoding, Hebei Province 071002, China; Beijing Key Laboratory of DNA Damage Response, College of Life Sciences, Capital Normal University, Beijing 100048, China; State Key Laboratory of Natural and Biomimetic Drugs, Chemical Biology Center, and School of Pharmaceutical Sciences, Peking University, Beijing 100191, China; CAS Key Laboratory of Genomic and Precision Medicine, Collaborative Innovation Center of Genetics and Development, College of Future Technology, Beijing Institute of Genomics, Chinese Academy of Sciences, 100101 Beijing, China; University of Chinese Academy of Sciences, 100049 Beijing, China; Institute of Stem Cell and Regeneration, Chinese Academy of Sciences, 100101 Beijing, China; Laboratory of Chemical Biology, Changchun Institute of Applied Chemistry, Chinese Academy of Sciences, Changchun, 130022, China

**Keywords:** O-GlcNAc, YTHDC1, m^6^A RNA, homologous recombination, DNA damage repair

## Abstract

N6-methyladenosine (m^6^A) is the most prevalent RNA modification, and its regulators include writers, readers and erasers. m^6^A is under stringent control and takes part in many biological events, but it is not known whether there is an interplay between m^6^A and glycosylation. Here we investigated an m^6^A reader, YTHDC1, which has been shown to be recruited to the DNA-RNA hybrid at DNA damage sites and regulate homologous recombination (HR) during DNA damage repair. We found that YTHDC1 is subject to O-linked β-N-acetylglucosamine (O-GlcNAc) modification at Ser396 upon DNA damage, which is pivotal for YTHDC1 chromatin binding and ionization radiation induced foci (IRIF) formation. RNA immunoprecipitation (RIP) and molecular dynamics (MD) simulations indicate that O-GlcNAcylation is vital for YTHDC1 to bind with m^6^A RNA. Fluorescence recovery after photo bleaching (FRAP) analysis revealed that YTHDC1 O-GlcNAcylation is essential for DNA damage-induced YTHDC1-m^6^A condensate formation. We further demonstrate that YTHDC1 O-GlcNAcylation promotes HR-mediated DNA damage repair and cell survival, probably through recruitment of Rad51 to the damage sites. We propose that YTHDC1 O-GlcNAcylation is instrumental for HR and genome stability.

## Introduction

O-linked β-N-acetylglucosamine (O-GlcNAc) is a dynamic glycosylation that occurs intracellularly (1). It is installed onto the Ser/Thr residues by the only writer O-GlcNAc transferase (OGT), and removed by the sole eraser O-GlcNAcase (OGA) (2). Currently, this quintessential post-translational modification (PTM) is found to decorate 5 000 proteins in the cytosol, nucleus and mitochondria and regulates various biological processes (3).

Emerging evidence suggests that the OGT and OGA pair play an important role in the DNA damage response (DDR) (4). Genotoxic stimuli from both the environment and our human body will induce various types of DNA damage, among which is DNA double-strand break (DSB) (5,6). DDR is the countermeasure against these lesions and in the case of DSB, it is repaired either by nonhomologous end joining (NHEJ) or homologous recombination (HR). NHEJ is more error-prone as it involves the ligation of DSB ends, and HR is more precise as its repair templates are the sister chromatids (7,8).

DDR induces protein O-GlcNAcylation and OGT is recruited to DNA damage sites (9). O-GlcNAcylation antagonizes DSB-induced phosphorylation of H2AX (γH2AX) to restrict γH2AX to the damage sites (9). OGA also relocates to DNA damage sites, albeit at a slower kinetics (9), which is mediated by the C-terminus of OGA (10). Substrates of OGT in DDR have been identified, including mediator of DNA damage checkpoint 1 (MDC1) (9), H2AX (9), and Non-POU Domain Containing Octamer Binding (NONO) (10) and Ku70/80 (10) (the latter two are in the NHEJ pathway). Apart from DSB repair, DNA polymerase eta (Polη) is O-GlcNAcylated during translesion DNA synthesis (TLS), which is a DDR pathway to respond to ultraviolet (UV)- and cisplatin-induced DNA lesions (11). During DNA synthesis, flap endonuclease 1 (FEN1) is O-GlcNAcylated and the abrogation of which will lead to DNA damage accumulation (12). Furthermore, High mobility group B1 (HMGB1) is O-GlcNAcylated, which alters its DNA binding ability and promotes an error-prone DNA repair (13). Recently, WD repeat and HMG-box DNA-binding protein 1 (WDHD1/AND-1) is also shown to be O-GlcNAcylated and takes part in HR (14). These data suggest that OGT and its resultant O-GlcNAcylation take part in various DDR pathways and are essential for genome integrity.

Here we demonstrate that YT521-B homology-domain-containing protein 1 (YTHDC1) O-GlcNAcylation is essential for HR. YTHDC1 is a reader of the N6-methyladenosine (m^6^A) mRNA. Initially discovered in 1974, m^6^A has been recently demonstrated to be the most prevalent mRNA modification (15,16). Writers and erasers of m^6^A have been identified, and its readers include the YTH family proteins (YTHDF1-3, and YTHDC1-2) (17). YTHDC1 has many biological roles (18,19). In RNA biology, it regulates mRNA splicing by binding with the pre-mRNA splicing factor SRSF3 (20). The YTHDC1-SRSF3 interaction could be disrupted by the Aurora A-dependent hnRNP K recruitment, resulting in exon skipping (21). YTHDC1 also promotes m^6^A mRNA translocation from the nucleus to the cytosol, and facilitates RNA-SRSF3 binding (22). In liquid-liquid phase separation (LLPS), YTHDC1-m^6^A forms condensates in acute myeloid leukemia (AML) cell lines, and YTHDC1 m^6^A-binding mutants display puncta formation and cell proliferation defects (23). In addition, m^6^A-modified enhancer RNA (eRNA) also binds with YTHDC1 to form condensates to promote gene activation (24).

YTHDC1 and other m^6^A factors take part in DDR. Upon DSB induction, the m^6^A writer Methyltransferase-like 3 (METTL3) is phosphorylated and relocated to the damage sites, where it mediates m^6^A modification in RNAs involved in DNA damage (25) (26). The resultant m^6^A further recruits YTHDC1 to promote DNA-RNA hybrid accumulation for subsequent HR-mediated DSB repair (26). *In vitro* studies suggest that the m^6^A writer complex Mettl3-MettL14 is located at UV-induced cyclopyrimidine dimer to promote correct DDR, and YTHDC1 is also caught at the scene of the crime(27). In breast cancer cells, Mettl3 promotes EGF expression via m^6^A and resultant YTHDC1 binding, leading to enhanced Rad51 expression and HR (28). Subsequently, the ablation of Mettl3 would impair HR and lead to Adriamycin (ADR) chemo-sensitivity in breast cancer cells (28). In AML cells and mouse models, YTHDC1 is essential for not only cell survival and proliferation, but also leukemogenesis, via the essential DNA replication helicase complex component minichromosome maintenance 4 (MCM4) (29).

In this paper we present evidence that YTHDC1 is O-GlcNAcylated at Ser396 during DNA damage. The glycosylation event promotes YTHDC1 chromatin binding and ionization radiation-induced focus (IRIF) formation. Through RNA immunoprecipitation (RIP) and molecular dynamics (MD) analysis, we show that O-GlcNAcylation promotes DNA damage-induced YTHDC1 condensate formation, probably through YTHDC1-m^6^A binding.

We further demonstrate that abrogation of the YTHDC1-m^6^A complex leads to γH2AX chromatin-dissociation defects. O-GlcNAcylated YTHDC1 enhances Rad51 recruitment and HR. Together, our results indicate that the OGT-YTHDC1-Rad51 axis promotes HR to protect genome integrity.

## Results

### DNA damage induces YTHDC1 O-GlcNAcylation at Ser396

Although O-GlcNAcylation has been known to play a role in DDR and it is elevated upon DNA damage (9), identification of its substrates has been hampered by its low stoichiometry, high lability and technical difficulties encountered in Mass Spec (MS) analysis (2). Recently a triaryl phosphine-trimethylpiperidine (TFT) reagent was developed to enable one-step identification of ionizing radiation (IR)-induced OGT substrates, and among the proteins identified is YTHDC1 (30). We therefore first examined the interaction between OGT and YTHDC1. Cells were transfected with HA-OGT and SFB-YTHDC1, and the cell lysates were immunoprecipitated (IPed) with anti-HA antibodies. And SFB-YTHDC1 was found to be in the immunoprecipitates (Fig. 1A). Reciprocally, OGT was also found in the SFB-YTHDC1 immunoprecipitates (Fig. 1B). Then pull-down assays were utilized. Cells were transfected with SFB-YTHDC1, and the cell lysates were incubated with recombinant GST-OGT proteins. GST-OGT could pulldown SFB-YTHDC1 (Fig. 1C). When endogenous interaction was examined, YTHDC1 proteins also coIP with OGT (Fig. 1D). These results suggest that YTHDC1 and OGT form a complex.

**Figure 1.**
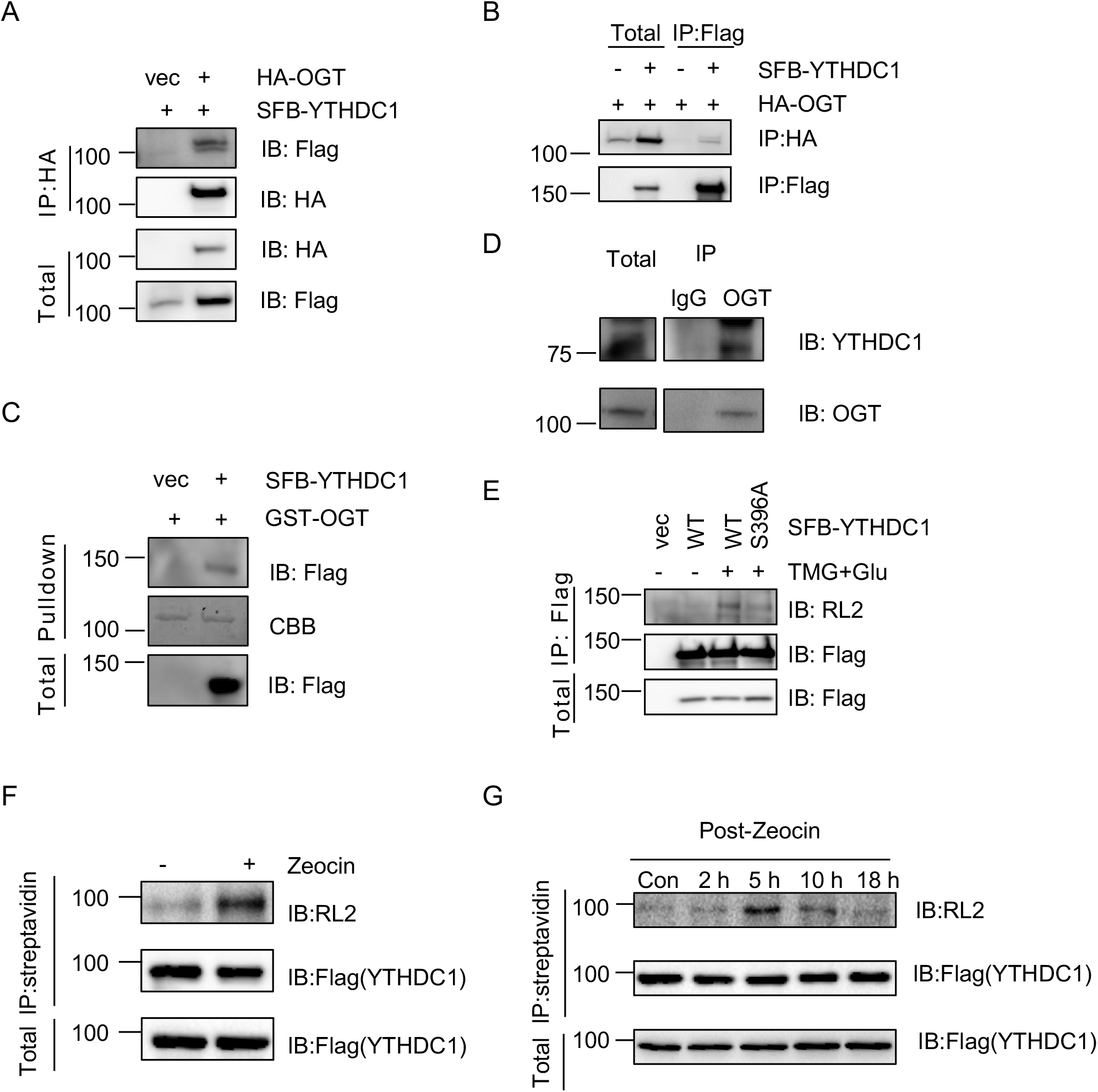
YTHDC1 is O-GlcNAcylated at S396 upon DNA damage. (A-B) Cells were transfected with HA-OGT and SFB-YTHDC1 plasmids. The cellular lysates were immunoprecipitated with anti-HA (A) or anti-Flag (B) antibodies and immunoblotted with the antibodies indicated. (C) Recombinant GST-OGT proteins could pulldown YTHDC1. Cells were transfected with SFB-YTHDC1 or vector controls. The cellular lysates were incubated with GST-OGT proteins purified from *E. coli*. (D) Endogenous OGT and YTHDC1 coIP. Cellular lysates were immunoprecipitated with anti-OGT antibodies and immunoblotted with anti-YTHDC1 antibodies. (E) Cells were transfected with YTHDC1-WT, or -S396A plasmids, treated or untreated with TMG plus glucose (31). The anti-Flag immunoprecipitates were subject to immunoprecipitation and immunoblotting with the antibodies indicated. (F) Zeocin (DNA damage agents) induces O-GlcNAcylation of YTHDC1. HCT116 cells were exposed to the DNA damaging agent Zeocin or mock treatment. The chromatin fractions were extracted and then incubated with streptavidin-conjugated beads at 4 °C for 2 h. The bound protein was subjected to SDS-PAGE and Western blotting with an anti-O-GlcNAc antibody (RL2). (G) DNA damage induces O-GlcNAcylation of YTHDC1 in a time course assay. HCT116 cells were exposed to 100 μg/ml of Zeocin for 4 hours and recovered for different time durations. The cell lysates were examined by Western blotting with anti-O-GlcNAc antibody (RL2) in a time course assay.

As the TFT-based method suggests that IR-induced O-GlcNAcylation occurs at Ser396, we generated S396A mutants of YTHDC1 and examined them for O-GlcNAcylation (Fig. 1E). Thiamet-G (TMG, an OGA inhibitor) and glucose incubation were utilized to enhance O-GlcNAcylation signals (31). S396A significantly diminished O-GlcNAcylation levels, suggesting that this is the main glycosylation site. We also assessed whether the modification responds to DNA damage by Zeocin treatment. As Fig. 1F shows, Zeocin incubation discernably increased YTHDC1 O-GlcNAcylation signals. When a Zeocin treatment time course was monitored (Fig. 1G), the O-GlcNAcylation of YTHDC1 peaked at approximately 5 hours, and then slowly started to recrease, indicative of a dynamic process of glycosylation.

### YTHDC1 O-GlcNAcylation Promotes Chromatin Binding and IR-induced focus (IRIF) formation

As YTHDC1 is recruited to DNA damage sites by m^6^A (26), we assessed the chromatin binding of YTHDC1-S396A mutants. As shown in Fig. 2A, YTHDC1-WT and -S396A mutants were transfected into cells and chromatin fractionation was carried out. We also treated the cells with Zeocin, a DNA damaging agent. As a result, S396A abolished the chromatin binding of YTHDC1. We also constructed GFP-YTHDC1 to visualize YTHDC1 localization in living cells. Under basal conditions, GFP-YTHDC1-WT and -S396A both form foci. Upon Zeocin treatment or IR, WT cells showed a robust increase of focus formation, while the S396A mutant was still at the basal level (Fig. 2B-C). We then examined γH2AX induction by cytology as it is a marker for DSB (26). In WT cells, Zeocin treatment significantly induced γH2AX localization, with a good correlation with GFP-YTHDC1 signals (Fig. 2D-E), consistent with previous observations (26). In S396A cells, however, γH2AX intensity correlated poorly with YTHDC1(Fig. 2D-E), suggesting that YTHDC1 O-GlcNAcylation is essential for its relocation to the damage sites.

**Figure 2.**
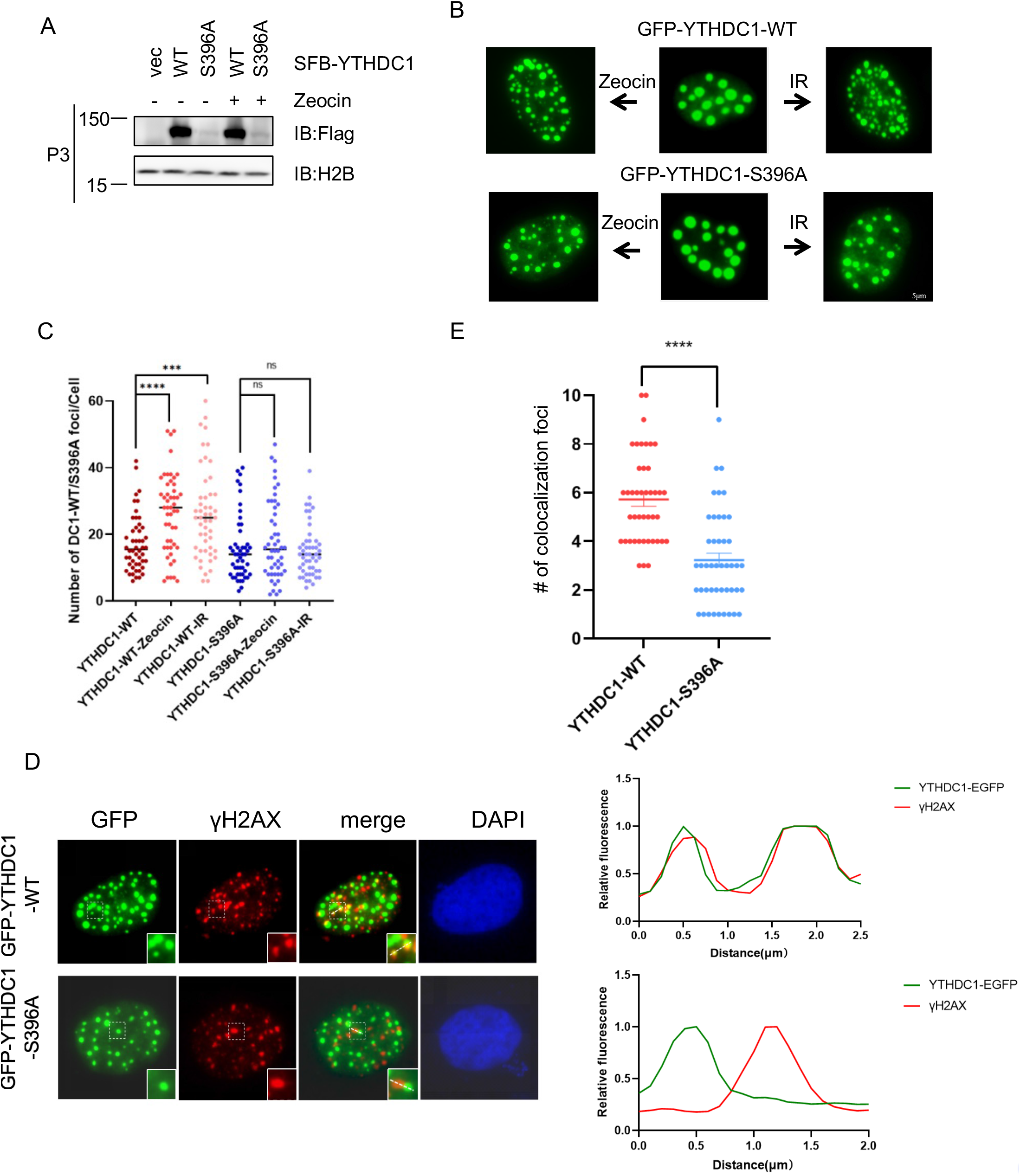
YTHDC1-S396A is defective in chromatin binding and ionizing radiation-induced focus (IRIF) formation. (A) Cells were transfected with YTHDC1-WT or -S396A plasmids and treated with Zeocin. The lysates were then subject to chromatin fractionation assays. The chromatin bound (P3) fraction was immunoblotted with anti-Flag antibodies to detect YTHDC1. (B) Cells were transfected with GFP-YTHDC1-WT, or -S396A plasmids, and then treated with 100 mg/ml Zeocin for 4 hrs, or 10 Gy X-ray. The GFP foci were quantitated in (C). 45 cells were counted in each experiment. ns: not significant; *** indicates P<0.001, **** indicates P<0.0001 (D) Cells were transfected with GFP-YTHDC1-WT, or -S396A plasmids, then treated with 10 Gy X-ray. The cells were then stained with anti-γH2AX antibodies, and the number of colocalization foci were quantitated in (E). **** indicates: P<0.0001

### YTHDC1 O-GlcNAcylation enhances m^6^A binding

As YTHDC1 is recruited to the damage sites via m^6^A binding(26), we tested the m^6^A binding ability of YTHDC1 via RNA immunoprecipitation (RIP) assays followed by dot blot (Fig. 3A-B). The S396A mutant significantly decreased m^6^A binding (Fig. 3A-B), which could be part of the reason for YTHDC1 localization defects during DDR.

**Figure 3.**
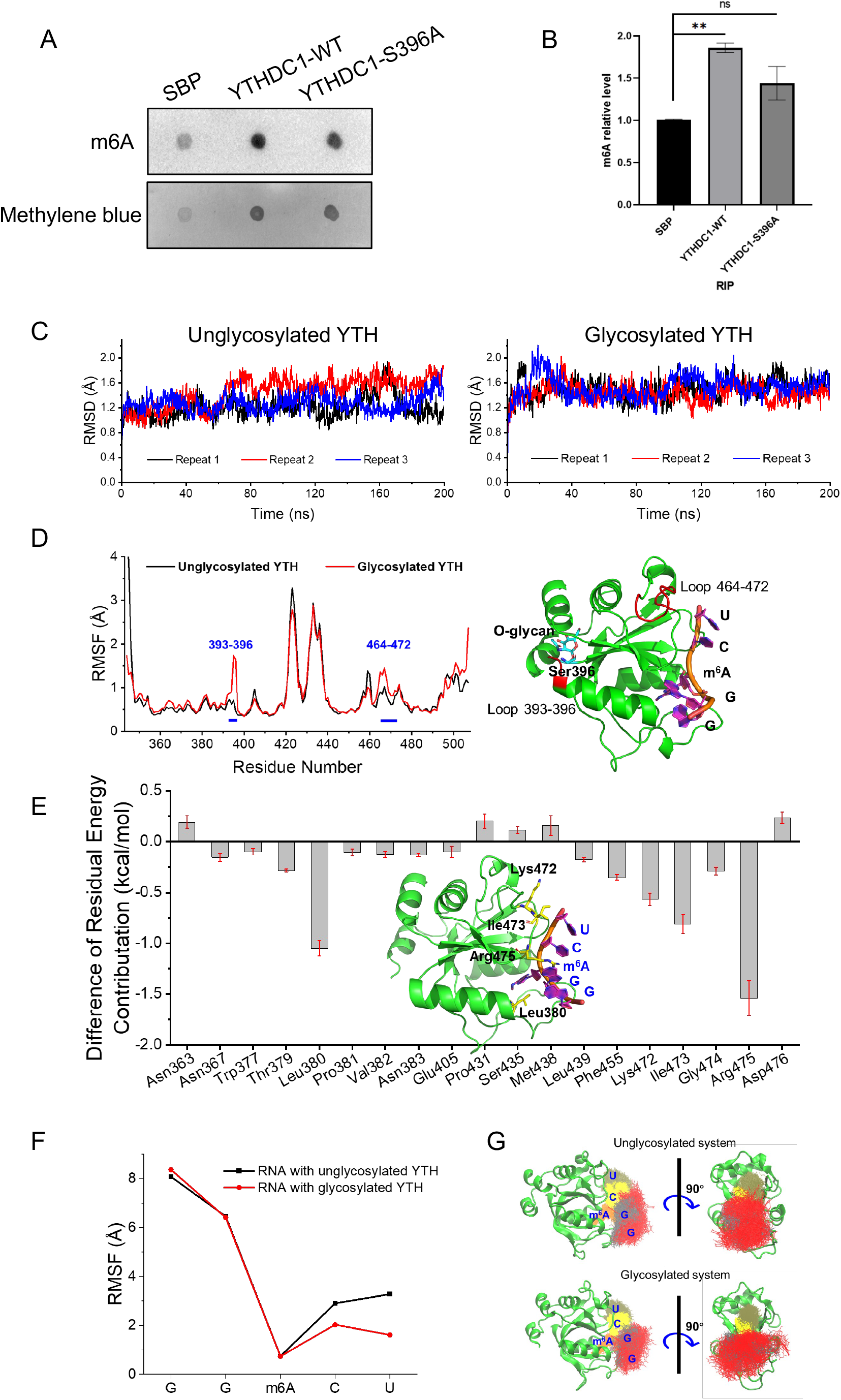
YTHDC1-S396A is defective in m^6^A mRNA binding. (A) RNA Immunoprecipitation (RIP) assay showed that YTHDC1-S396A is defective in m^6^A mRNA binding. (B) Quantitation of (A). ns indicates not significant, ** indicates p<0.01. (C-G) Molecular dynamics simulations indicate that glycosylated YTHDC1 reduced m^6^A mRNA binding through an allosteric regulation. (C) Root-mean-square deviations (RMSDs) of Cα atoms of unglycosylated YTH (left) and glycosylated YTH (right) with RNA in the independent repeated simulations (Repeat 1, black; Repeat 2, red; and Repeat 3, blue). (D) Root mean square fluctuations (RMSFs) of the Cα atoms of unglycosylated (black) and glycosylated (red) YTH in the combined 240 ns trajectories. (E) Differences in each residue energy contribution between the unglycosylated system and the glycosylated system (only the residues with contributions > 0.1 kcal/mol are shown). The inset shows a schematic view of the key residues around RNA. (F) RMSF of bases in unglycosylated (black) and glycosylated (red) YTH in the combined 240 ns trajectories. (G) Overlay of RNA structures from the combined 240 ns trajectories.

Aiming to probe the potential modulatory role of O-GlcNAc on Ser396 at the molecular level, we first attempted to synthesize the O-GlcNAcylated YTH domain of YTHDC1 via a chemical approach, which should afford a homogeneous glycoprotein that would be inaccessible through bioexpression protocols. Accordingly, we dissected the YTH domain into four segments 1a-4a that could be prepared using solid-phase peptide synthesis and were subsequently assembled based on chemical ligation and desulfurization strategies (Figure S1-2). While the peptidyl hydrazide-based ligations proceeded efficiently, we noticed that the desulfurization steps were problematic, either leading to incomplete conversion of intermediates, such as peptide 6a, or generating insoluble aggregates at the final stage of assembling full length protein 10a. Eventually, we managed to obtain a small amount of the S396-GlcNAcylated YTH domain (11a). The synthetic protein displays an analogous CD spectrum to that of the nonglycosylated YTH domain obtained from *E. coli* expression (Figure S3), suggesting comparable secondary structures. Unfortunately, the synthetic route was unable to provide sufficient materials for further biochemical and biological evaluations.

To circumvent the difficulties of accessing S396-GlcNAcylated YTHDC1, we resorted molecular dynamics (MD) simulation. The root-mean-square deviation (RMSD) values were first calculated. As shown in Fig 3C, both systems can reach stable states in 100 ns. The frames of the last 80 ns were extracted from the trajectory of each repeat and combined together for further analysis for each system (240 ns in total). To verify whether O-GlcNAc influences the fluctuation of each residue during simulations, the root-mean-square fluctuation (RMSFs) were calculated for the combined 240 ns trajectories. The existence of O-GlcNAc not only disturbed the loop (Residue 393-396) it is located, but also increased the fluctuation of the loop close to the RNA binding site (Residue 464-472) (Fig 3D). The binding energy of glycosylated YTH to RNA was -89.29 ± 0.48 kcal/mol, which is lower than that of unglycosylated YTH to RNA (−78.93 ± 0.54 kcal/mol). The differences in each residue contribution between the unglycosylated system and the glycosylated system are shown in Fig 3E. After S396 was O-GlcNAcylated, RNA GG(m^6^A)CU was further stabilized by Leu380 and Ile473 with van der Waals interactions and Lys472 and Arg475 with electrostatic interactions.

The behavior of RNA in the binding site was also investigated. The methylated adenosine (m^6^A) deeply inserted the binding site and was not influenced by O-glycosylation (Fig 3F). Regardless of whether S396 was O-GlcNAcylated, the GG motif of RNA was equally flexible. However, glycosylation dramatically restricted the fluctuation of the CU motif of RNA (Fig 3G). Overall, the glycosylation increased the binding affinity of RNA to the YTH domain and stabilized the binding of RNA, probably through an allosteric regulation.

### YTHDC1 O-GlcNAcylation facilitates DNA damage-induced YTHDC1-m^6^A condensate formation

Recent work suggests that O-GlcNAcylation inhibits LLPS (32,33), and YTHDC1-m^6^A is implicated in LLPS (23) (24), we therefore examined the effect on condensate formation by YTHDC1. As shown in Fig 4A-B, fluorescence recovery after photo bleaching (FRAP) assays indicated that ∼50% GFP-YTHDC1-S396A recovered ∼50 seconds after photobleaching, but WT displayed much slower kinetics and a much lower intensity, suggesting that O-GlcNAcylation is indeed against LLPS under basal conditions.

**Figure 4.**
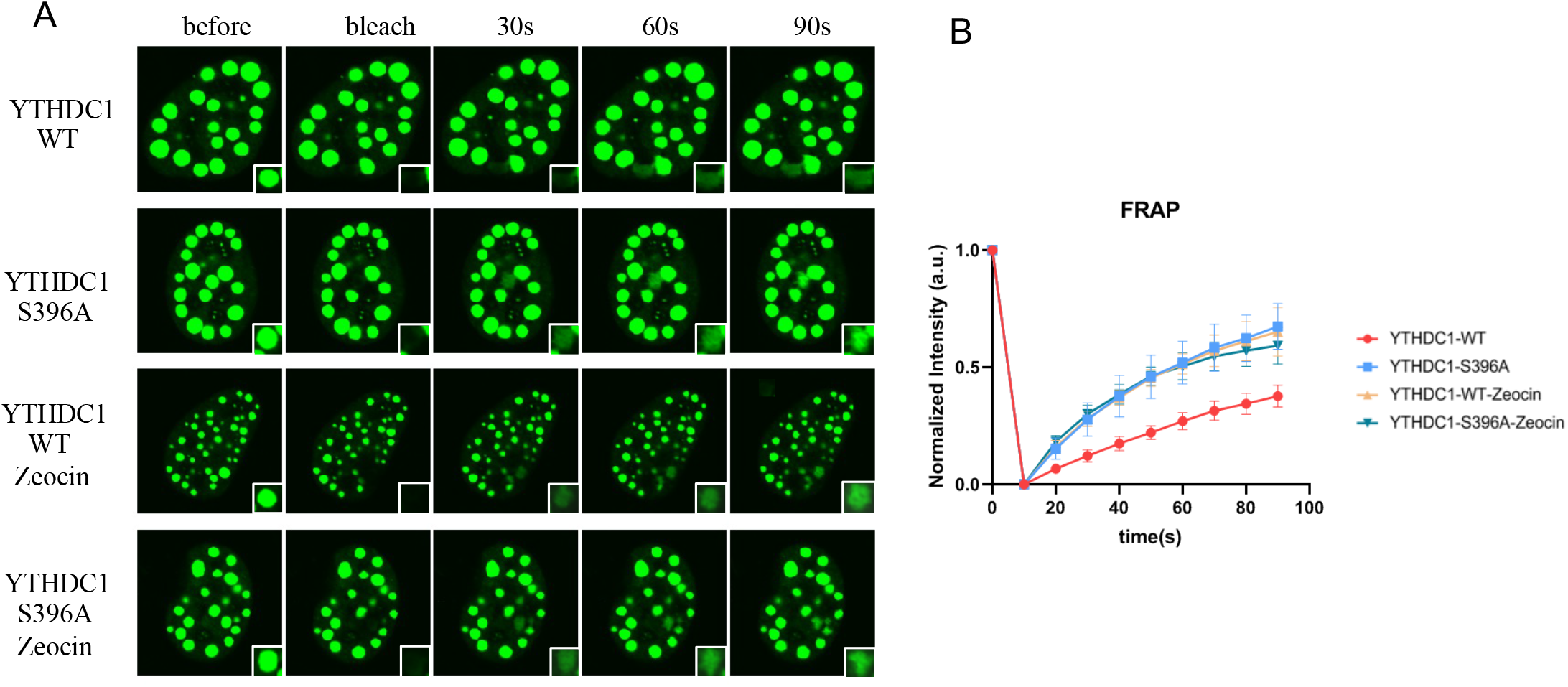
YTHDC1 O-GlcNAcylation enhances YTHDC1-m^6^A condensate formation upon DNA damage. (A) Fluorescence recovery after photo bleaching (FRAP) of GFP-YTHDC1-WT and -S396A droplets *in vivo*,with or without Zeocin treatment. Scale bar, 5 µM. (B) Fluorescence intensity recovery of YTHDC1 droplets after photobleaching (N=3).

Upon Zeocin treatment, however, YTHDC1-WT and S396A responded completely differently. YTHDC1-WT manifested much more DNA damage-induced foci, consistent with Fig. 2B. And Zeocin treatment increased the LLPS rate in YTHDC1-WT-transfected cells, consistent with the recent finding that m^6^A could enhance the phase separation of many m^6^A reader proteins (34-36), including YTHDC1 (23). The FRAP rate in YTHDC1-S396A, however, remained the same as that in untreated cells, indicating that DNA damage-induced m^6^A could account for the difference.

### YTHDC1 O-GlcNAcylation promotes HR

Since the data above suggest a role for YTHDC1 O-GlcNAcylation in regulating DDR, we sought to determine whether it regulates HR or NHEJ. Using the established GFP reporter assays (37), we found that YTHDC1 modulates HR (Fig. 5A), but not NHEJ (Fig. 5B), consistent with published results (26,28). We then transfected YTHDC1-S396A into the DR-GFP system to monitor HR, and the results showed decreased HR efficiency (Fig. 5C), suggesting that S396 O-GlcNAcylation is essential for HR.

**Figure 5.**
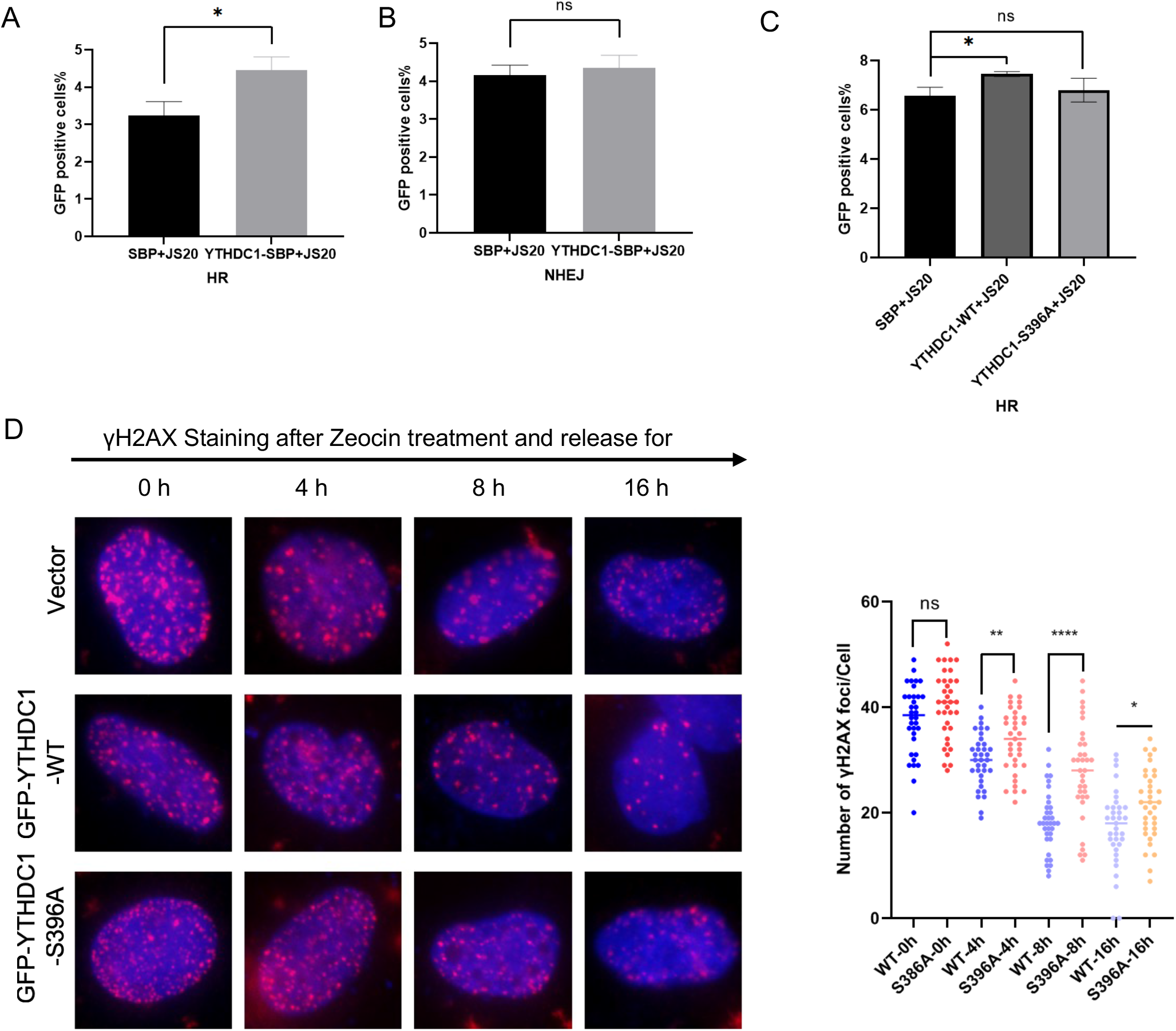
YTHDC1 O-GlcNAcylation promotes DSB repair. (A) DR-U2OS cells were infected with I-SceI and YTHDC1-WT or control plasmids to detect homologous recombination (HR). The percentage of GFP-positive cells is shown. YTHDC1-WT transfected cells showed more HR-mediated DSB repair. * indicates P<0.05. (B) EJ5-GFP 293 cells were transfected with I-SceI and YTHDC1-WT or control plasmids to detect non-homologous end joining (NHEJ). The percentage of GFP-positive cells is shown. There is no significant difference between YTHDC1-WT and control cells. NS: not significant. (C) DR-U2OS cells were infected with I-SceI and YTHDC1-WT or -S396A plasmids. HR-mediated DSB repair was measured by flow cytometry. NS: not significant. * indicates P<0.05. (D-E) YTHDC1-S396A cells displayed defects in γH2AX dissociation. Cells were transfected with GFP-YTHDC1-WT, or -S396A plasmids, treated with Zeocin and released for the time indicated. Then the cells were stained with anti-γH2AX antibodies and quantitated (E). Scale bar, 5 µM. ns: not significant; * indicates P<0.05, ** indicates P<0.01, **** indicates P<0.0001.

We next analyzed _γ_H2AX dissociation after Zeocin treatment, as it is an indication of DDR completion (38). As shown in Fig. 5D, _γ_H2AX foci started to decrease rapidly 4 hours after release from Zeocin treatment, and continued to decline until 16 hours. In contrast, in the S396A cells, _γ_H2AX foci remained much more abundant compared to the WT. This is consistent with an HR defect in the S396A mutant.

### YTHDC1 O-GlcNAcylation promotes HR through Rad51 recruitment

Then we sought to identify the downstream targets of YTHDC1. We first depleted endogenous YTHDC1 with siRNA, and then transfected YTHDC1-WT and -S396A (Fig. 6A). Using these cells, we then measured for Rad51 recruitment, because the METTL3-m^6^A-YTHDC1 axis is implicated in Rad51 recruitment (26). As shown in Fig. 6B-C, siYTHDC1 depletion significantly reduced Rad51 focus formation, in line with previous findings that YTHDC1 upregulates Rad51 expression (28). Compared with YTHDC1-WT, S396A significantly attenuated Rad51 focus formation (Fig. 6B-C), suggesting that YTHDC1 O-GlcNAcylation is essential for Rad51 localization to DSB sites.

**Figure 6.**
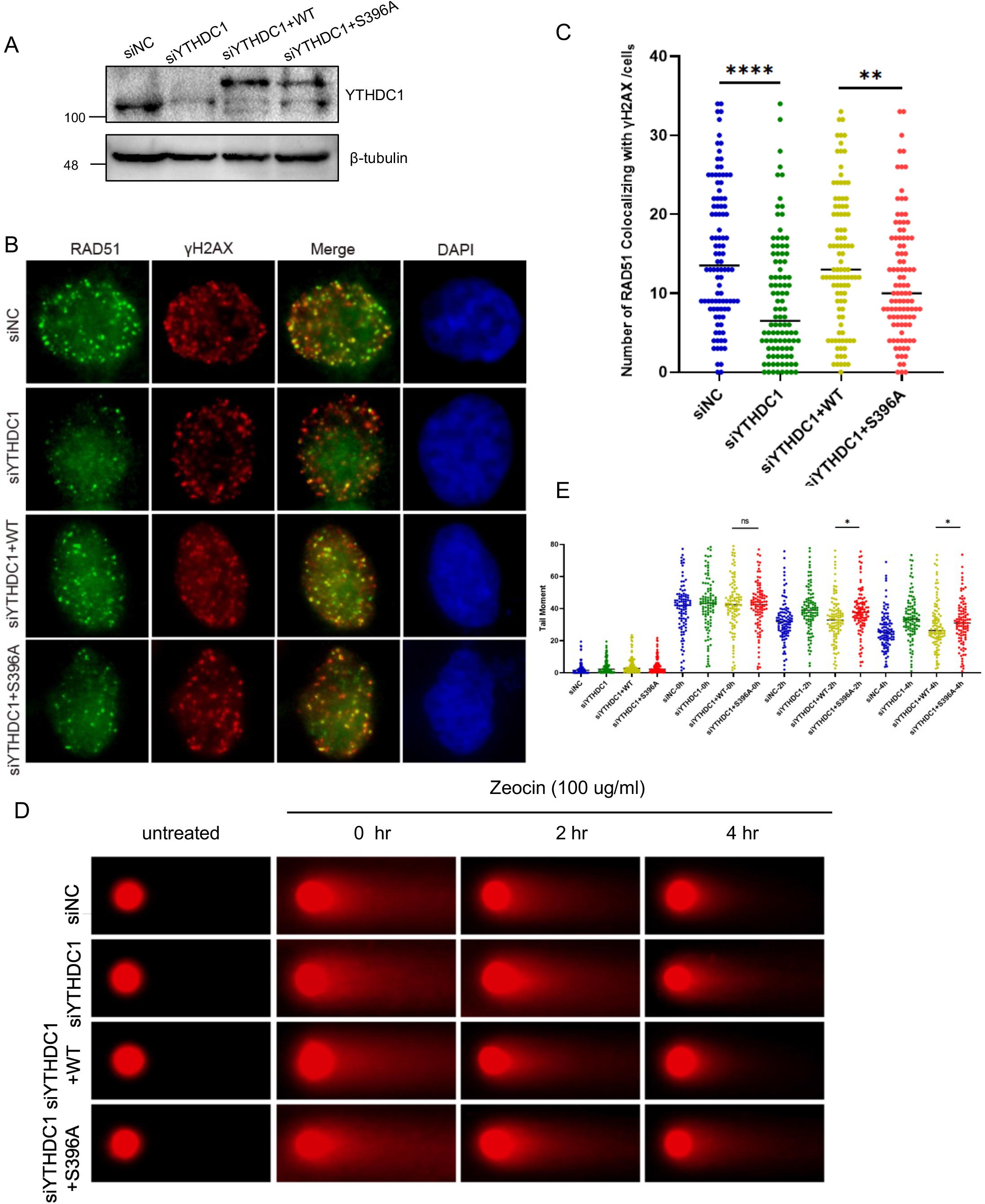
YTHDC1-WT modulates HR through Rad51 recruitment. (A) Cells were treated with siRNA targeting YTHDC1 and then transfected with YTHDC1-WT or S396A plasmids. (B) The cells in (A) were treated with Zeocin, and co-stained with anti-Rad51 and anti-γH2AX antibodies. (C) Quantitation of the co-localization between Rad51 and γH2AX. ** indicates P<0.01, **** indicates P<0.0001. (D) YTHDC1-S396A is defective in DNA damage repair as analyzed by neutral comet assays. The cells in (A) were treated with Zeocin and then recovered for the time indicated. Comet tails were analyzed, and the DNA damage repair kinetics were assessed (E). ns indicates not significant, * indicates p<0.05.

Then using the same cells, we measured for DSB repair kinetics by comet assays (10). Zeocin was used to induce DSBs, and knockdown of YTHDC1 or reconstitution of S396A significantly impaired DSB repair (Fig. 6D-E). Furthermore, we utilized different doses of Zeocin to treat the cells, and carried out colony formation assays (Fig. 7A-B). Again, the S396A mutant displayed hypersensitivity. Collectively, these results suggest that YTHDC1 O-GlcNAcylation is important for DDR, perhaps through Rad51 recruitment.

**Figure 7.**
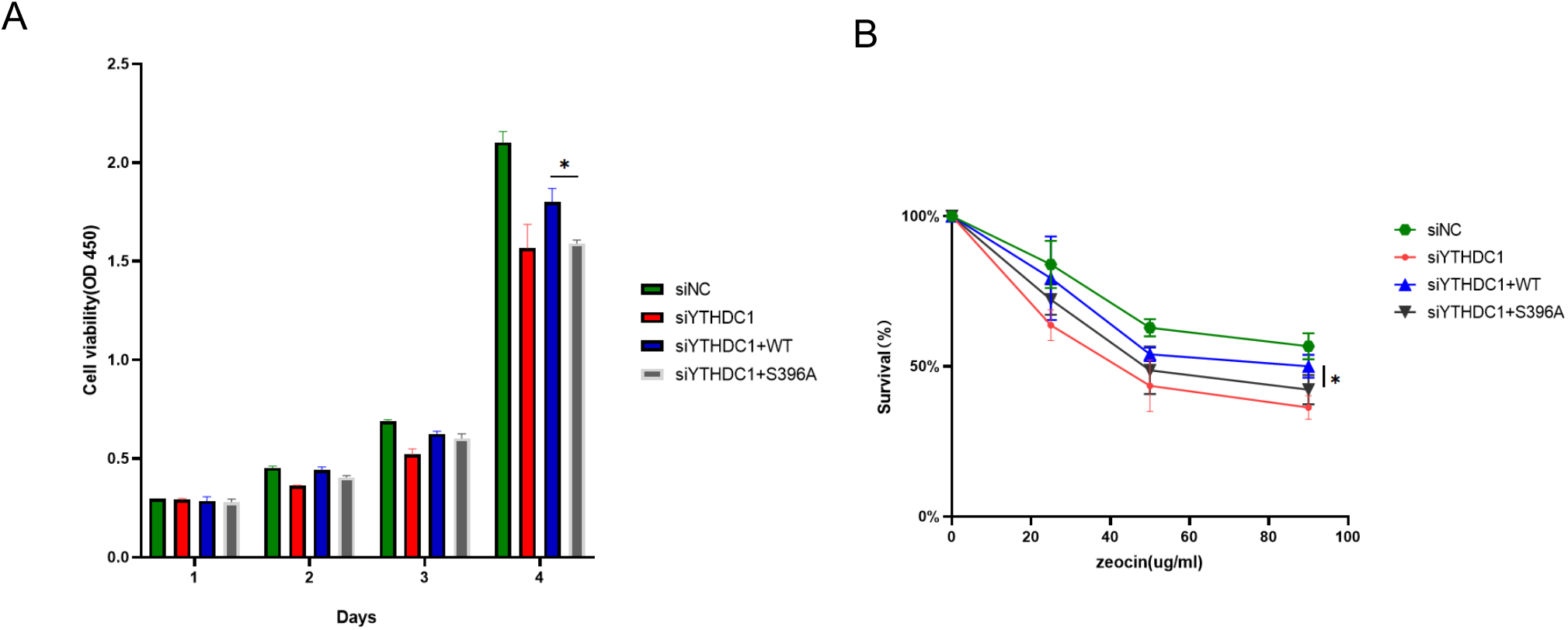
YTHDC1 O-GlcNAcylation is essential for cell survival. The cells were treated with Zeocin for different durations or different doses of Zeocin, and colony formation assays were performed in the indicated cell lines. * indicates p<0.05.

## Discussion

In this work, we present evidence that YTHDC1, an m^6^A reader, is O-GlcNAcylation upon DNA damage. This glycosylation event is indispensable for YTHDC1 chromatin-binding, m^6^A RNA binding and consequent recruitment to DNA damage sites. Through biochemical and cytological analysis, we show that YTHDC1 O-GlcNAcylation promotes _γ_H2AX recruitment and Rad51 localization and enhances DDR through HR. Recently small RNAs are shown to be N-glycosylated on the cell surface (39), our work demonstrates that the m^6^A mRNA reader YTHDC1 is O-GlcNAcylated in the cell, suggesting that the interconnection between RNA and glycosylation could be more than meets the eye.

Among the OGT substrates that have been identified, the O-GlcNAcylated sites often reside in intrinsically disordered regions (IDRs). On the contrary, YTHDC1 O-GlcNAcylation occurs at S396, right on the YTH domain, which is the m^6^A binding domain. Through RIP assays and MD, we show that S396 O-GlcNAcylation is important for YTHDC1-m^6^A binding. We think there might be other O-GlcNAcylated sites on functional domains, in addition to IDRs.

Recently, the relationship between O-GlcNAc and LLPS has been under intense investigation. For instance, neuronal SynGAP T1306 O-GlcNAc has been demonstrated to inhibit the formation of the SynGAP-PSD-95 complex, and thus LLPS (33). In addition, O-GlcNAc would attenuate LLPS of the RNA-binding EWS N-terminal disordered region (32). In our work, under basal conditions, we also found that O-GlcNAcylation inhibited YTHDC1 LLPS (Fig. 4A). However, upon DNA damage, YTHDC1-S396A displayed no significant changes compared to the untreated control. YTHDC1-WT, however, started to manifest phase separation phenotypes, probably due to increased binding with m^6^A and thus formation of the YTHDC1-m^6^A condensates. These results suggest that the effect of O-GlcNAcylation on LLPS differs in distinct cellular contexts, and that different binding partners might exert various effects on LLPS.

Structure-based MD simulations have gained popularity to facilitate the functional probing of glycosylated proteins, probably due to the current difficulty in the chemical synthesis of glycopeptides or glycoproteins. Previously MD has been employed to predict glycoengineering of GPCRs with O-GlcNAc, and the simulation suggests that the modification would enhance proteolytic stability (40). A recent work on malate dehydrogenase 1 (MDH1) with MD indicates that O-GlcNAc strengthens MDH1-substrate binding, probably as a molecular glue (41). When chemical synthesis does succeed, it no doubt pinpoints the exact role of O-GlcNAc on the specific protein or peptide. For instance, when a removable glycosylation modification (RGM) method was used in total chemical synthesis, O-GlcNAc was found to improve disulfide-rich protein folding (42). In the neurodegenerative disease-causing _α_-synuclein, O-GlcNAc inhibits amyloid aggregation (43). In our case, although we could not synthesize the O-GlcNAcylated YTHDC1 protein, our MD results suggest that O-GlcNAc could impose an allosteric regulation of YTHDC1, thus enhancing YTHDC1-m^6^A binding. This is yet another unappreciated function of O-GlcNAcylation. In the future, perhaps a combination of chemical synthesis and MD would generate more interesting results and reveal more multifaceted roles of O-GlcNAc.

The exact function of OGT in DDR is murky waters. Due to the difficulty in pinpointing OGT modification sites, it is not yet clear whether OGT or OGA plays a major role in HR or NHEJ. Both OGT and OGA are recruited to the damage sites, although OGA is recruited at a slower speed (9). It is a possible scenario that OGT gets on the DNA damage sites first to catalyze protein O-GlcNAcylation, and after the substrates have carried out their function, it is OGA’s turn to reverse the reaction. A second scenario is that these two enzymes catalyze distinct sets of substrates for totally different functions. Reports of O-GlcNAcylated proteins in both pathways have been anecdotal: MDC1(9), H2AX (9), AND-1(14) in HR, and NONO (10) and Ku70/80 (10) in NHEJ. The potential role of O-GlcNAc in other types of damage, such as DNA single-strand break, has not been investigated. Due to the transient nature of DNA damage and the repair process, new methodologies have been developed to identify the O-GlcNAcylated substrates in a time-resolved fashion (12,30). Perhaps a more concerted role of OGT or OGA in DDR would be unveiled with more chemoproteomic studies in the future.

## Materials and methods

### Cells and antibodies

Cells were grown under standard conditions. The following plasmids were described: YTHDC1 plasmids (20), and OGT plasmids (44). YTHDC1-S396A was generated following the manufacturers’ instructions (QuickChange II, Stratagene). The primary antibodies were as follows: YTHDC1 (Abcam, ab 122340), _γ_H2AX (Abcam, ab26350), Rad51 (Abcam, ab133534), and m^6^A (Synaptic Systems, 202 003). Chemicals: TMG plus glucose (TMG + Glu) (31) treatment was TMG (5 _μ_M) for 24 h and 30 mM glucose for 3 hours; Zeocin: 100 _μ_g/mL for 4 h. siRNA sequences are as follows:

siYTHDC1-1: GTCGACCAGAAGATTATGATA

siYTHDC1-2: ATCGAGTATGCAAATATTGAA

### Cell lysis, immunoprecipitation (IP), and Western Blotting

The cells were harvested at the indicated time points after relevant treatment and washed twice with phosphate-buffered saline (PBS). The cell pellets were subsequently resuspended in the NETN lysis buffer (20 mM Tris-HCl, pH 8.0, 300 mM NaCl, 1 mM EDTA and 0.5 % NP-40). For IB, the lysates were used with indicated antibodies and incubated with Pierce Protein G Agarose (Thermo) for 2–4 h at 4 □. After washing the agarose beads with NETN100 buffers (20 mM Tris-HCl, pH 8.0, 100 mM NaCl, 1 mM EDTA and 1% NP-40), the samples were mixed with protein loading buffers at 100 □ for 5 min. The protein lysates were separated using 10% SDS □ polyacrylamide gel electrophoresis and equal quantities (5 µg per sample) of separated proteins were transferred to 0.22 □ µm nitrocellulose membranes (MilliporeSigma). After blocking with 5% non□fat milk for 1 h at room temperature, the membranes were incubated with specific primary antibodies overnight at 4°C. After washing three times with PBS, the membranes were incubated with secondary antibodies for 2 h at room temperature. The proteins were visualized using an enhanced chemiluminescence detection kit (Amersham; Cytiva) and the Odyssey Infrared Imaging system version 2.1 (LI □ COR Biosciences).

### DNA damage-induced focus formation assay

Cells were cultured on coverslips and treated with 100 μg/mL Zeocin (a radio-mimetic chemical that induces DSBs) for 4 h. After recovery for 4 h, cells were fixed in 4% paraformaldehyde at room temperature for 15 min, and permeabilized with 0.5% Triton X-100 in PBS for 10 min. Samples were blocked with blocking buffer (8% goat serum in PBS) and then incubated with anti-γH2AX antibodies, followed by fluorescence-labeled secondary antibodies for 1 h. The nuclei were stained with DAPI. The number of foci was counted in at least 100 cells/sample.

### RNA immunoprecipitation (RIP) assay

Control and YTHDC1-WT or -S396A overexpressed cells were collected and washed twice with ice-cold PBS. Cells were lysed in ice-cold IP lysis buffers (20 mM Tris-HCl, pH 8.0, 300 mM NaCl, 1 mM EDTA and 0.5 % NP-40) for 30 min on ice and spun down at maximum speed to precipitate the debris at 4 □. Supernatants were collected and incubated with 5 ug anti-YTHDC1 antibodies or an equivalent amount of rabbit IgG by rotating overnight at 4□. RNA-YTHDC1-antibody complexes were pulled down using Dynabeads Protein A/G (Millipore) and washed 3 times with NETN100 buffers (20 mM Tris-HCl, pH 8.0, 100 mM NaCl, 1 mM EDTA and 1% NP-40). Proteinase K (Thermo Fisher Scientific Inc.) was applied to remove proteins from the complexes.

### Molecular dynamics (MD) dimulations

The crystal structure of the YTHDC1 YTH domain with m^6^A RNA was extracted from the complex of RNA GG(m^6^A)CU and YTH (PDB ID: 4R3I) (45). The O-glycan (β-*N*-Acetyl-_D_-Glucosamine) at S396 of YTHDC1 was built using the Glycan Reader & Modeler module(46).

The role of O-glycosylation in YTHDC1 recognition of RNA was investigated via molecular dynamics simulations. Two systems (unglycosylated YTHDC1 and O-GlcNAcylated YTHDC1 at S396 in complex with RNA, respectively) solvated in TIP3P water molecules with 150 mM KCl were built by the CHARMM-GUI webserver(47). The simulations were performed by the Gromacs 2021.2 program (48,49)with the CHARMM36m force field (50,51). The systems were equilibrated in the isothermal-isobaric (NPT) ensemble at a temperature of 310 K for 200 ns. The SHAKE algorithm was applied to restrain all bonds involving hydrogen (52). The particle-mesh Ewald (PME) summation method was applied to treat long-range electrostatic interactions (53). The pressure was set at 1 atm maintained by the Nosé–Hoover Langevin piston method (54). Each system was repeated three times independently.

The RMSD (root-mean-square deviation) and RMSF (root-mean-square fluctuation) analyses were performed through MDAnalysis (55). The binding energy (enthalpy) and per-residue energy contributions were calculated by the molecular mechanics/Poisson-Boltzmann (generalized-Born) surface area method with the gmx_MMPBSA tool (56,57). The interactions between YTH and RNA were displayed by PyMol (58).

### Fluorescence recovery after photo bleaching (FRAP)

For FRAP experiments in living cells, an area of 1 mm-diameter of YTHDC1 puncta was bleached with a 405 nm laser using an Olympus FV3000 Laser-Scanning Confocal Spectral Inverted Microscope. The GFP fluorescence signal was collected over time. Each data point is representative of the mean and standard deviation of fluorescence intensities in three unbleached (control) or three bleached (experimental) granules. The pre-bleached fluorescence intensity was normalized to 1 and the signal after bleach was normalized to the prebleach level.

### Homologous recombination (HR)/non-homologous end joining (NHEJ) *in vivo* reporter assays

DR-GFP-U2OS cells and EJ5-GFP-U2OS cells were overexpressed with YTHDC1-WT or YTHDC1-S396A, and then transfected with JS-20 (SceI) or empty vectors. After 48 h, 2 × 10^5^ cells were analyzed for GFP-positive cells by flow cytometry to demonstrate the repair efficiency of HR and NHEJ. In flow cytometry figures, the y-axis is side scattered light (SSC) to determine the granularity of cells. The x-axis is the GFP signal intensity of cells.

GFP-positive populations were gated to distinguish a group of cells with positive signals.

Uninduced cells were included as negative controls. The results represent the mean value of triplicate experiments.

### Comet assays

To evaluat DNA DSBs, single-cell gel electrophoretic comet assays were performed under neutral conditions. HCT116 cells were overexpressed with YTHDC1-WT or YTHDC1-S396A plasmids with or without 100 μg/mL Zeocin. After incubating in fresh medium for the indicated time at 37 □, the cells were harvested and mixed with 1% LMP agarose at 42 □ and immediately pipetted onto slides. To perform neutral conditional assays, the slides were immersed in the neutral lysis buffer (2% sarkosyl, 0.5 M EDTA, 0.5 mg/mL proteinase K, pH 8.0) overnight at 4 □. After lysis, the slides were washed with the electrophoresis buffer (90 mM Tris-HCl at pH 8.5, 90 mM boric acid, 2 mM EDTA), and analyzed by electrophoresis at 25 V for 40 min (0.6 V/cm) and stained in 10 μg/mL propidium iodide for 30 min in the dark. All images were taken under a fluorescence microscope and analyzed by the Comet Assay IV software program.

## Data Availability Statement

All data are contained in this manuscript.

## Abbreviations

(FRAP): Fluorescence recovery after photo bleaching
(HR): homologous recombination
(NHEJ): non-homologous end joining
(m^6^A): N6-methyladenosine
(IP): immunoprecipitation
(IB): immunoblotting
(DSB): Double-strand break
(RIP): RNA Immunoprecipitation
(O-GlcNAc): O-linked β-N-acetylglucosamine
(OGT): O-GlcNAc transferase
(YTH): YT521-B homology
(YTHDC1): YTH Domain Containing 1
(MD): Molecular Dynamics

## Acknowledgement

Jing L. is supported by the National Natural Science Foundation of China (NSFC) fund (31872720) and R & D Program of Beijing Municipal Education Commission (KZ202210028043). C. W. is supported by NSFC (32071277), Natural Science Foundation of Hebei province (C2021201012), S&T Program of Hebei (216Z2602G) and Interdisciplinary Research Program of Natural Science of Hebei University (DXK202006).

