## Supplemental figures for "DNA damage-induced YTHDC1 O-GlcNAcylation promotes homologous recombination by enhancing N6-methyladenosine binding"

**Materials and Methods**

HPLC grade tetrahydrofuran, methylene chloride, diethyl ether, toluene, and N, N-Dimethylformamide were purchased from Fisher, Acros, J&K, and Oceanpak, and were purified and dried by passing through a PURE SOLV® solvent purification system (Innovative Technology, Inc.) when anhydrous solvents were required. Other reagent grade solvents for chromatography were purchased from Beijing Tongguang Fine Chemicals Company. Molecular sieves (4 Å, power) were pre-activated in an oven at 65 °C overnight and further flame-dried before being used in the reactions. All other reagents and relevant catalysts were purchased from Sigma-Aldrich, NovaBiochem, GL Biochem, Acros, TCI, Adamas, Innochem, J&K, Alfa, and Energy, and were used without further purification. Ultra-pure argon (≥99.999%) was used when inert reaction conditions were required. Ultraviolet-Visible Spectroscopy was performed on a WPA Biowave II Spectrophotometer. Analytical thin layer chromatography was performed using 0.25 mm silica gel 60-F plates (Merck). Flash chromatography was performed using 200-300 mesh silica gel (Qingdao Haiyang Chemical Co., Ltd.). Yields refer to chromatographically and spectroscopically pure materials unless otherwise stated. ^1^H, ^13^C, and two-dimensional NMR spectra were recorded on a Bruker Ultrashield Plus 400 MHz or 600 MHz spectrometer at ambient temperature. Data for ^1^H NMR are reported as follows: chemical shift, integration, multiplicity (ovrlp = overlapping, s = singlet, d = doublet, t = triplet, q = quartet, m = multiplet) and coupling constants. All ^13^C NMR spectra were recorded with complete proton decoupling.

All HPLC separations involved a mobile phase of 0.05% (v/v) TFA in water (solvent A) and 0.04% (v/v) TFA in MeCN (solvent B) unless otherwise stated. All peptide UV traces were recorded under the wavelength of 210 nm and 220 nm in analytical HPLC-MS, analytical HPLC and preparative HPLC.

Analytical HPLC-MS chromatographic separations were performed using a Waters Alliance e2695 Separations Module, an SQ Detector, and a Waters 2489 UV/Visible (UV/Vis) Detector equipped with an Exsil Pure 300 C18 column (5.0 μm, 4.6 × 150 mm) at a flow rate of 0.4 mL/min or a Higgins analytical Proto-300 C4 (5.0 μm, 2.1 × 150 mm) at a flow rate of 0.2 mL/min. All of the LC-MS/MS data were obtained from Thermo Xcalibur 2.2 SP1.48. The UV-Vis chromatograms of analytical LC-MS were acquired from Waters 2489 UV/Visible detector, and the MS spectra of analytical LC-MS were acquired from Waters SQD mass spectrometry. LC-MS data deconvolution was processed by MassLynx (version 4.1).

Preparative HPLC separations were performed using a Hanbon Sci. & Tech. NP7005C solvent delivery system and a Hanbon Sci. & Tech. NU3010C UV detector equipped with an Exsil Pure 300 C18 column (10 μm, 20 × 250 mm) at a flow rate of 15 mL/min or a Proto 300 C4 column (10.0 μm, 20 × 250 mm) at a flow rate of 16 mL/min.

Automated solid-phase peptide synthesis, Removal of acetyl protections of carbohydrate on peptide, Preparation of peptidyl acids and peptidyl hydrazides and manually coupling GlcNAcylated Ser were described as before ^[1]^.

**Hydrazide-based native chemical ligation**
Peptidyl hydrazide was dissolved to 1-5 mM in 6 M Gn·HCl with 200 mm MPAA (pH 3.0), 1 equiv acac (from a 150 mM stock in water) were added to the mixture, and the reaction mixture was stirred for 4 hours to form thioester. Cysteine-containing fragment (1.2 equiv) was dissolved in 6 M Gn·HCl with 200 mm Na_2_HPO_4_ at pH 8.5, and added to the mixture. This ligation mixture was brought to pH 7 with the addition of 1 M NaOH and allowed to sit overnight. The reaction was monitored by LC MS, and reducing buffer (6 M Gn·HCl, 200 mM Na_2_HPO_4_, 150 mM TCEP·HCl, pH 7.2) was added to the reaction solution and incubated for 20 minutes, and the reaction was then quenched by adding equal volume of CH_3_CN/H_2_O/AcOH (30/65/5, v/v/v) and further purified using HPLC.

**Metal-free desulfurization**

To a solution of the Cys-containing peptide in 200 μL of degassed buffer (6 M Gn·HCl, 200 mM Na_2_HPO_4_, pH 7.2) was added 200 μL of 0.5 M Bond-breaker^®^ TCEP solution (Pierce), 80 μL of 2-methyl-2-propanethiol and 100 μL of radical initiator VA-044 (0.1 M in degassed water). The reaction mixture was stirred at 37 °C for 5 hours, quenched by addition of CH_3_CN/H_2_O/AcOH (30/65/5, v/v/v) and further purified using HPLC.

**Removal of Acm groups on cysteines**

5 mg of PdCl_2_ was dissolved in 500 μL of degassed neutral buffer (6 M Gn·HCl, 200

mM Na_2_HPO_4_) under an argon atmosphere, and the mixture was gently stirred for 30

min at 37 °C for complete dissolution. To the solution of Acm-protected peptide in

neutral buffer was added the PdCl_2_ (10 equiv) solution, and the resulting mixture was

stirred at 37 °C for 30 min. Then 2-fold volume of degassed guanidine buffer (6 M

Gn·HCl, 1 M DTT) was added, and the resulting mixture was further incubated for 15

min. The solution was diluted with CH_3_CN/H_2_O/AcOH (30/65/5, v/v/v) and further

purified by HPLC

**Synthesis of Ser-GlcNAc**

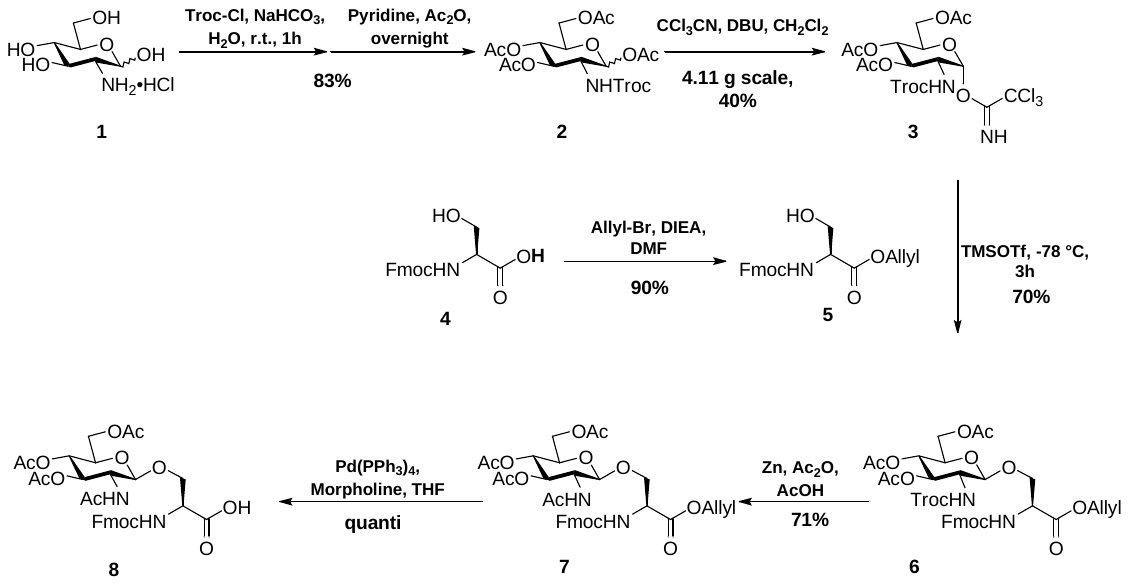

The synthetic route and characterization of compounds were described as before ^[2]^.

**Synthesis of YTHDC1-GlcNAc**

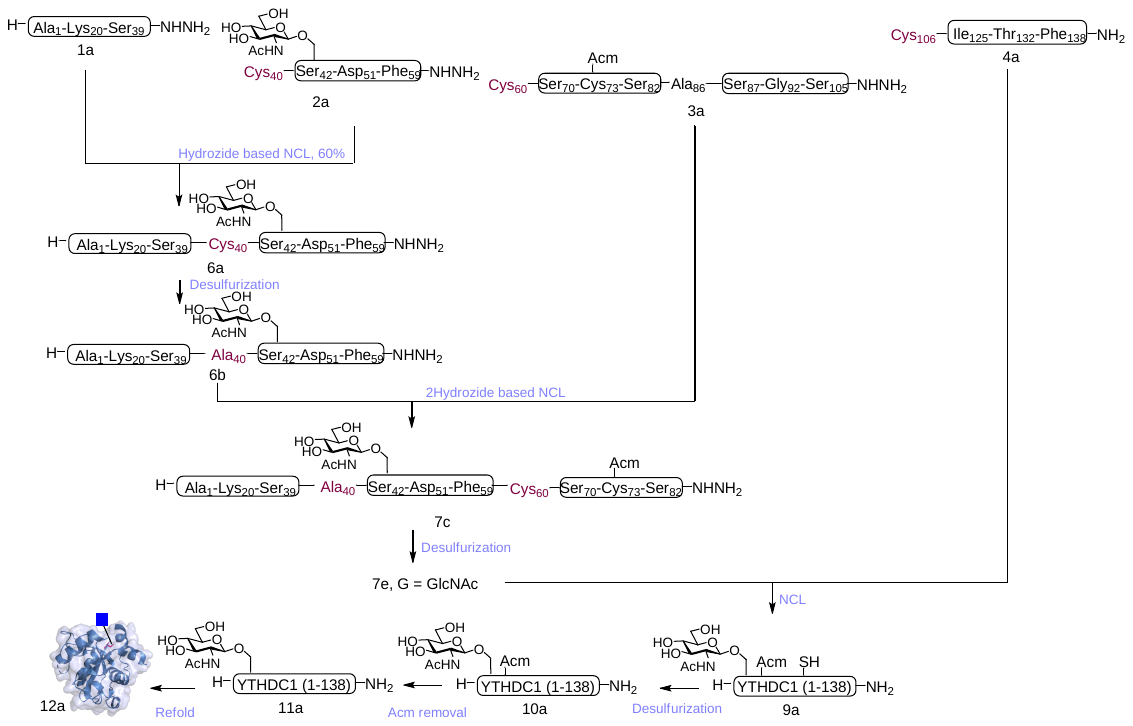

Peptidyl Hydrazide 1a

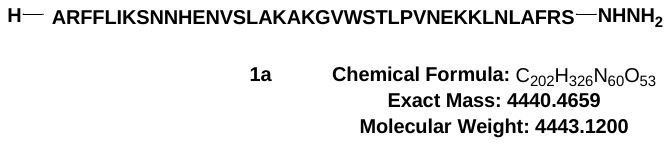

Peptide 1a was prepared according to General Procedure using Fmoc-NHNH-CTC resin (0.28 mmol/g, 0.05 mmol), After global deprotection using TFA/TIS/H_2_O (95/2.5/2.5, v/v/v), the crude peptide was dissolved in 10 mL of CH_3_CN/H_2_O/AcOH (35/60/5, v/v/v) and further purified using RP-HPLC (linear gradient 20-45% solvent B over 30 min, Proto-300 C4 column-1). Product was collected and lyophilized to provide peptide 1a (24 mg, 11%) as a white powder.

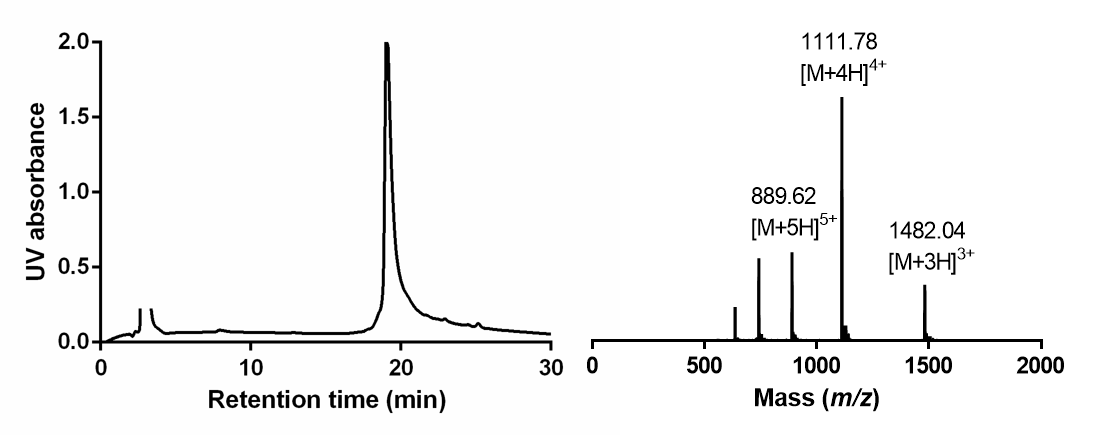

HPLC-MS analysis of **1a**. Left: UV traces (Liner gradient of 25-40% solvent B over 30 min, Proto-300 C4 column, t_R_ = 19.6 min); Right: ESI-MS data. Calcd mass for C_202_H_326_N_60_O_53_: 4443.12 Da (average isotopes), [M+3H]^3+^ m/z = 1482.04, [M+4H]^4+^ m/z =1111.78, [M+5H]^5+^ m/z =889.62; observed: 1482.04, 1111.78, 889.62.

Peptidyl Hydrazide 2a

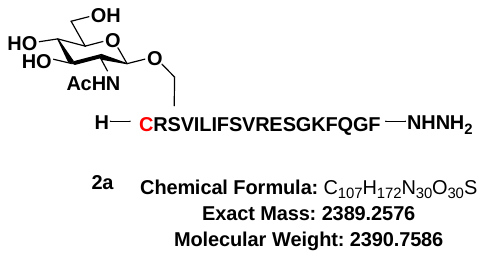

Peptide 2a was prepared according to General Procedure using Fmoc-NHNH-CTC resin (0.28 mmol/g, 0.05 mmol), Fmoc-Ser(Ac3GlcNAc)-OH (**8**) was coupled as previously described. After global deprotection using TFA/TIS/H_2_O (95/2.5/2.5, v/v/v), the crude peptide was dissolved in 10 mL of CH_3_CN/H_2_O/AcOH (35/60/5, v/v/v) and purified using RP-HPLC (linear gradient 10-40% solvent B over 30 min, Exsil Pure 300 C18 column). Product was collected and lyophilized to provide peptide 2a (32mg, 27%) as a white powder.

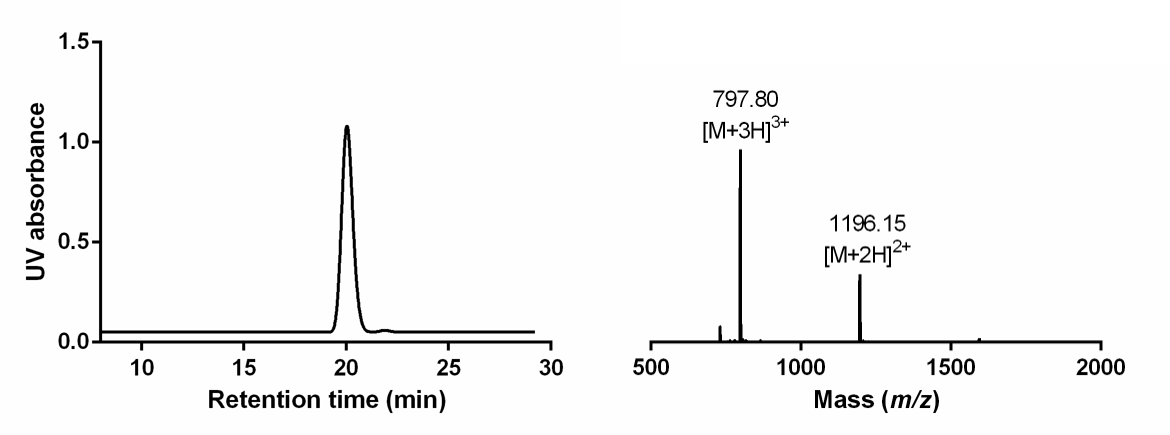

HPLC-MS analysis of **2a**. Left: UV traces (Liner gradient of 10-40% solvent B over 30 min, Exsil Pure 300 C18 column, t_R_ = 20.1 min); Right: ESI-MS data. Calcd mass for C_107_H_172_N_30_O_30_S: 2390.75 Da (average isotopes), [M+2H]^2+^ m/z = 1196.37, [M+3H]^3+^ m/z = 797.91; observed:1196.15, 797.80.

Peptidyl Hydrazide 3a

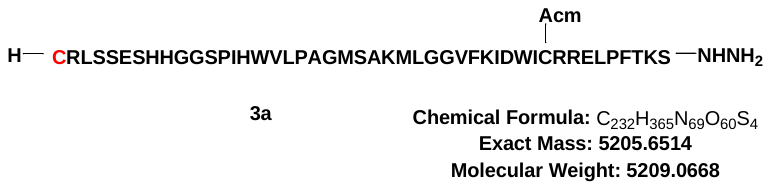

Peptide 3a was prepared according to General Procedure using Fmoc-NHNH-CTC resin (0.28 mmol/g, 0.05 mmol), After global deprotection using TFA/TIS/H_2_O (95/2.5/2.5, v/v/v), the crude peptide was dissolved in 10 mL of CH_3_CN/H_2_O/AcOH (35/60/5, v/v/v) and further purified using RP-HPLC (linear gradient 25-45% solvent B over 30 min, Proto-300 C4 column-1). Product was collected and lyophilized to provide peptide 3a (17mg, 6.5 %) as a white powder.

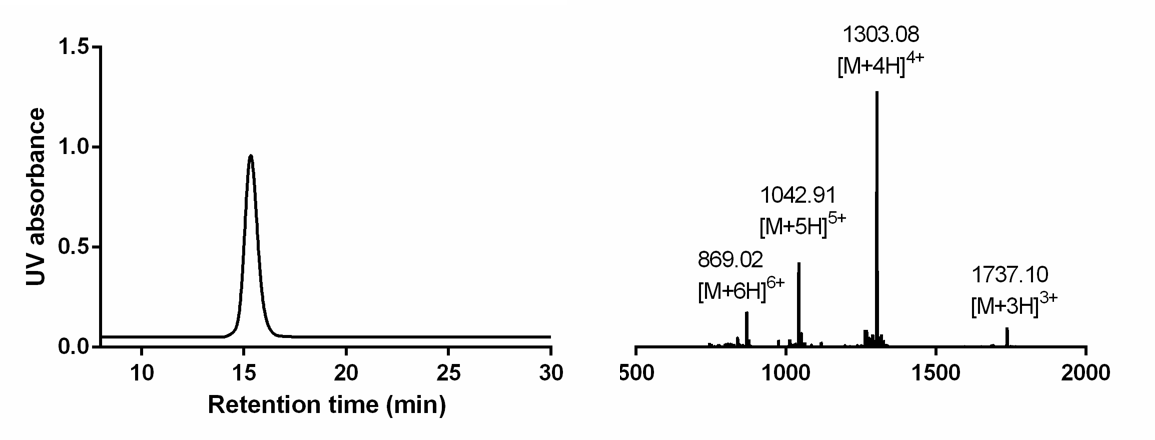

HPLC-MS analysis of **3a**. Left: UV traces (Liner gradient of 25-45% solvent B over 30 min, Proto-300 C4 column, t_R_ = 15.2 min); Right: ESI-MS data. Calcd mass for C_232_H_365_N_69_O_60_S_4_: 5209.06 Da (average isotopes), [M+3H]^3+^ m/z = 1737.35, [M+4H]^4+^ m/z = 1303.26, [M+5H]^5+^ m/z = 1042.81, [M+6H]^6+^ = 869.17; observed:1737.10, 1303.08, 1042.91, 869.02.

Peptidyl Hydrazide 4a

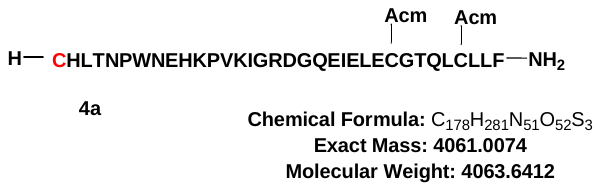

Peptide 4a was prepared according to General Procedure using Fmoc-Rink Amide MABA resin (0.37 mmol/g, 0.05 mmol), After global deprotection using TFA/TIS/H_2_O (95/2.5/2.5, v/v/v), the crude peptide was dissolved in 10 mL of CH_3_CN/H_2_O/AcOH (35/60/5, v/v/v) and further purified using RP-HPLC (linear gradient 25-45% solvent B over 30 min, Proto-300 C4 column-1). Product was collected and lyophilized to provide peptide 4a (24 mg, 11%) as a white powder.

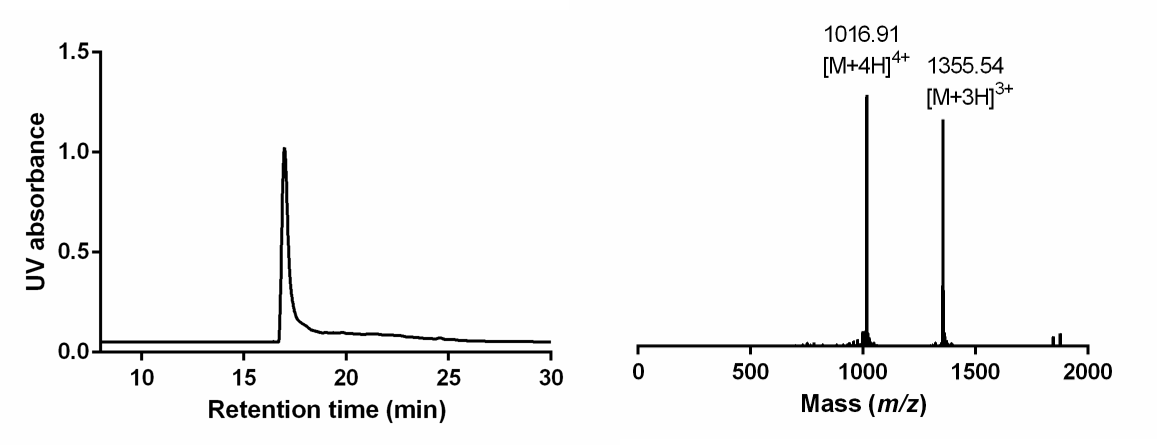

HPLC-MS analysis of **4a**. Left: UV traces (Liner gradient of 25-45% solvent B over 30 min, Proto-300 C4 column, t_R_ = 18.1 min); Right: ESI-MS data. Calcd mass for C_178_H_281_N_51_O_52_S_3_: 4063.64 Da (average isotopes), [M+3H]^3+^ m/z =1355.54, [M+4H]^4+^ m/z = 1016.91; observed: 1355.54, 1016.91.

Peptidyl Hydrazide 6a

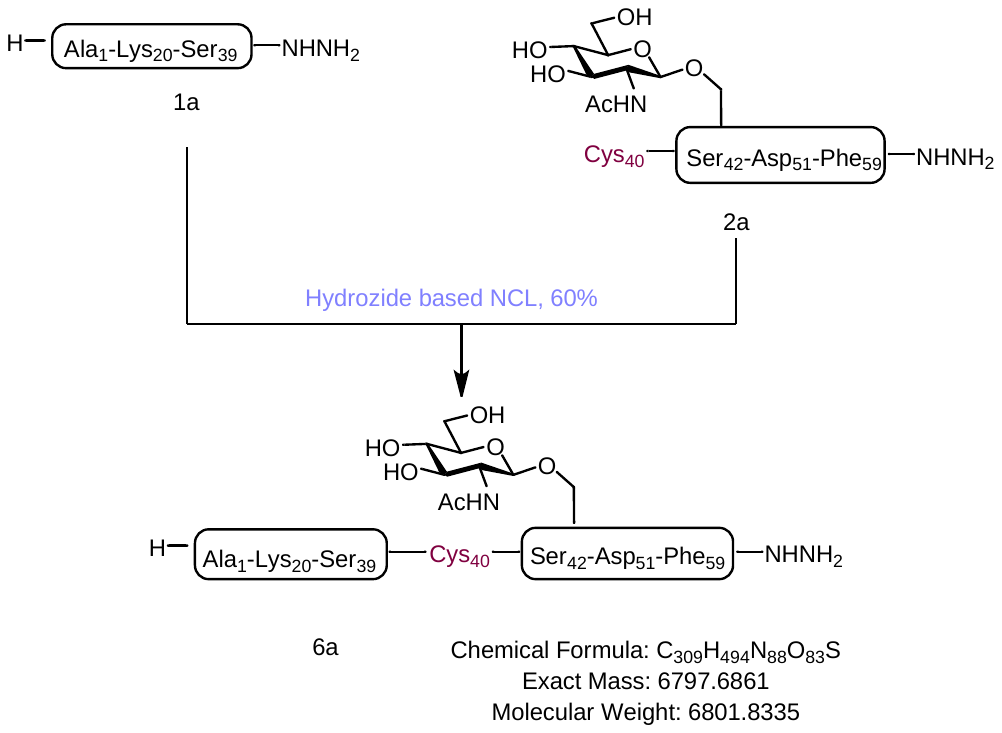

Peptidyl hydrazide 1a (4 mg, 0.90 µmol) and peptide 2a (2.26 mg, 0.90 µmol) were subjected to the ligation conditions following General Procedure as described previously. The reaction was stirred at room temperature for 12 hours, followed by addition of reducing buffer (6 M Gn·HCl, 200 mM Na_2_HPO_4_, 150 mM TCEP·HCl, pH 7.2) and quenched by CH_3_CN/H_2_O/AcOH (30/65/5, v/v/v). The ligated product was purified using RP-HPLC (linear gradient 25-45% solvent B over 30 min, Proto-300 C4 column-1). Product was collected and lyophilized to provide peptide 6a (4.43 mg, 72%) as a fluffy white solid.

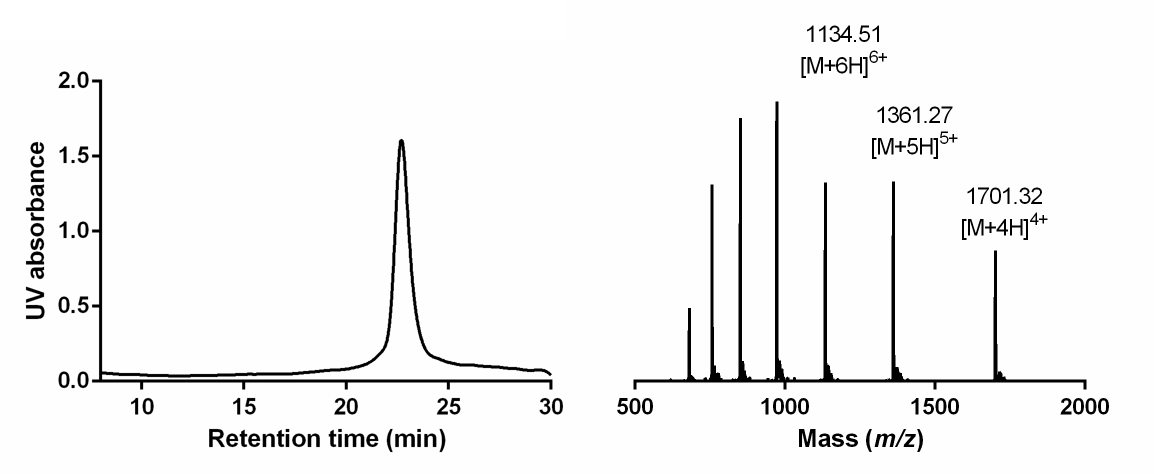

HPLC-MS analysis of **6a**. Left: UV traces (Liner gradient of 25-40% solvent B over 30 min, Proto-300 C4 column, t_R_ = 22.3 min); Right: ESI-MS data. Calcd mass for C_309_H_494_N_88_O_83_S: 6801.83 Da (average isotopes), [M+4H]^4+^ m/z =1701.45, [M+5H]^5+^ m/z =1361.36, [M+6H]^6+^ m/z =1134.63; observed: 1701.32, 1361.27, 1134.51.

Peptidyl Hydrazide 6b

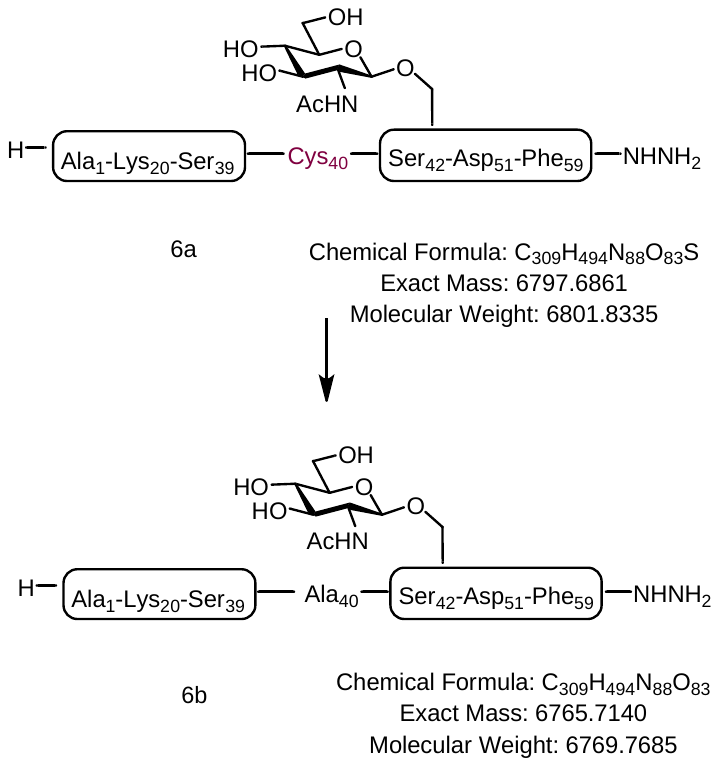

Peptide 6a (4.43 mg, 0.65 µmol) were subjected to the MFD conditions following General Procedure as described previously. The reaction was stirred at 37 ℃ for 5 hours, it need twice reaction, followed by addition of reducing buffer (6 M Gn·HCl, 200 mM Na_2_HPO_4_, 150 mM TCEP·HCl, pH 6.2) and quenched by CH_3_CN/H_2_O/AcOH (30/65/5, v/v/v). The product was purified using RP-HPLC (linear gradient 25-45% solvent B over 30 min, Proto-300 C4 column-1). Product was collected and lyophilized to provide peptide 6b (1.12 mg, 25%) as a fluffy white solid

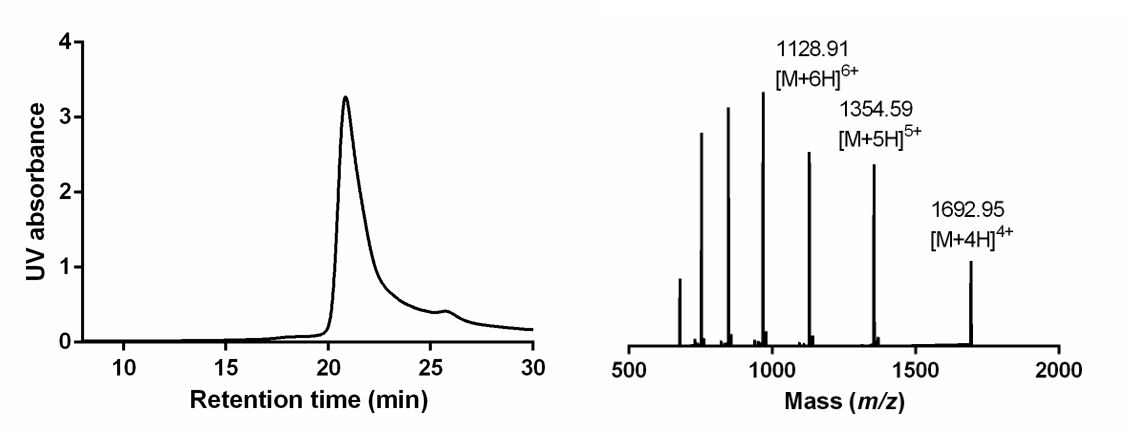

HPLC-MS analysis of **6b**. Left: UV traces (Liner gradient of 25-45% solvent B over 30 min, Proto-300 C4 column, t_R_ = 21.2 min); Right: ESI-MS data. Calcd mass for C_309_H_496_N_88_O_83_: 6769.76 Da (average isotopes), [M+4H]^4+^ m/z = 1693.44, [M+5H]^5+^ m/z = 1354.95, [M+6H]^6+^ m/z = 1129.29; observed: 1692.95, 1354.59, 1128.91.

Peptidyl Hydrazide 7a

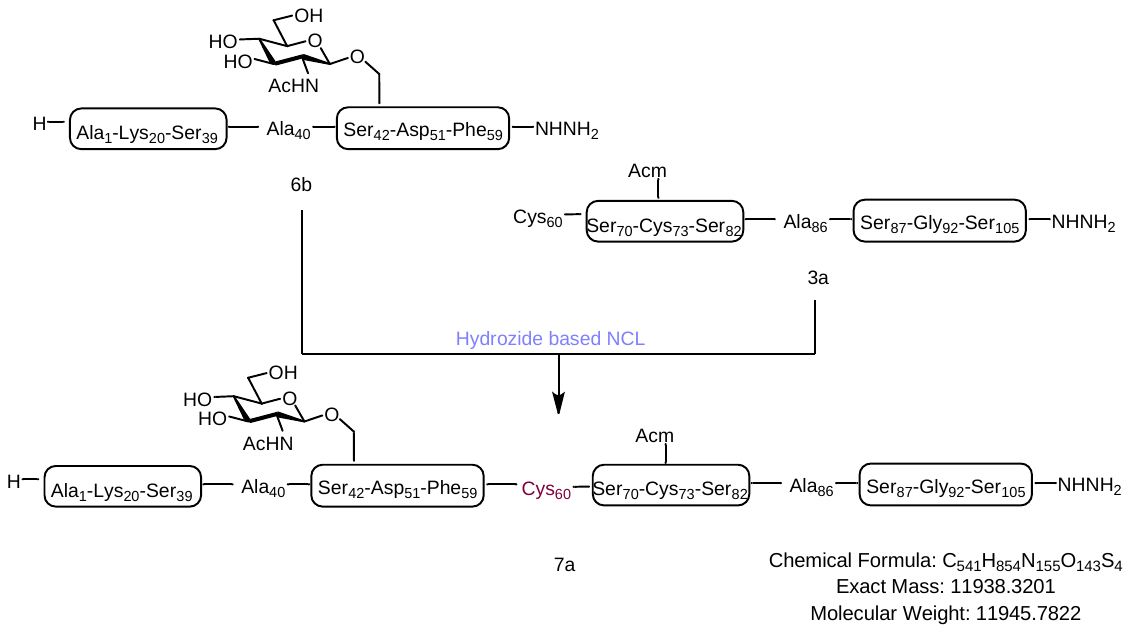

Peptidyl hydrazide 6b (2.59 mg, 0.38 µmol) and peptide 3a (2 mg, 0.38 µmol) were subjected to the ligation conditions following General Procedure as described previously. The reaction was stirred at room temperature for 12 hours, followed by addition of reducing buffer (6 M Gn·HCl, 200 mM Na_2_HPO_4_, 150 mM TCEP·HCl, pH 7.2) and quenched by CH_3_CN/H_2_O/AcOH (30/65/5, v/v/v). The ligated product was purified using RP-HPLC (linear gradient 30-45% solvent B over 30 min, Proto-300 C4 column-1). Product was collected and lyophilized to provide peptide 7a (1.13 mg, 25%) as a fluffy white solid.

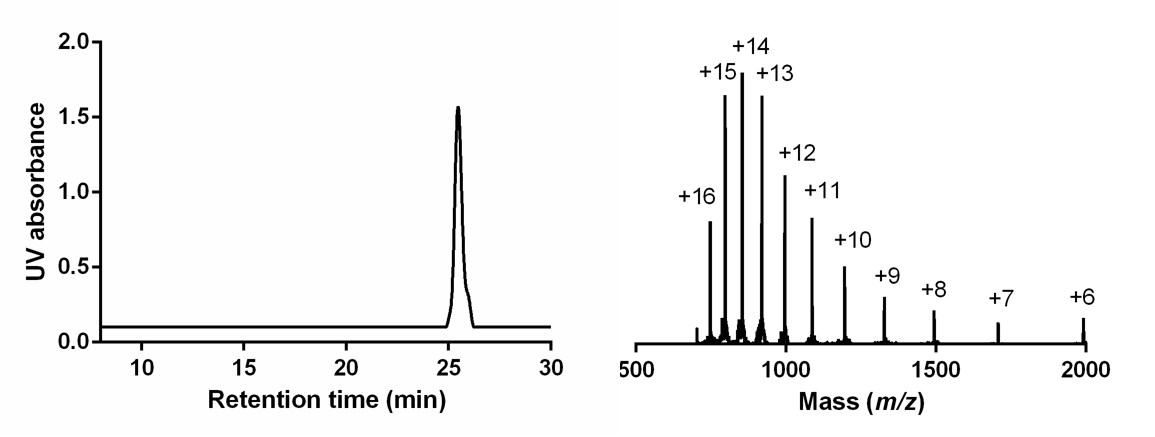

HPLC-MS analysis of **7a**. Left: UV traces (Liner gradient of 30-45% solvent B over 30 min, Proto-300 C4 column, t_R_ = 26.2 min); Right: ESI-MS data. Calcd mass for C_541_H_854_N_155_O_143_S_4_: 11945.78Da (average isotopes), [M+6H]^6+^ m/z =1991.96, [M+7H]^7+^ m/z =1707.54, [M+8H]^8+^ m/z =1494.22, [M+9H]^9+^ m/z =1328.30, [M+10H]^10+^ m/z =1195.57, [M+11H]^11+^ m/z =1086.98, [M+12H]^12+^ m/z =996.48, [M+13H]^13+^ m/z =919.90, [M+14H]^14+^ m/z =854.27, [M+15H]^15+^ m/z =797.38, [M+16H]^16+^ m/z = 747.61; observed: 1990.99, 1706.79, 1493.87, 1327.93, 1195.34, 1086.72, 996.27, 919.65, 854.10, 797.20, 747.45.

Peptidyl Hydrazide 7b

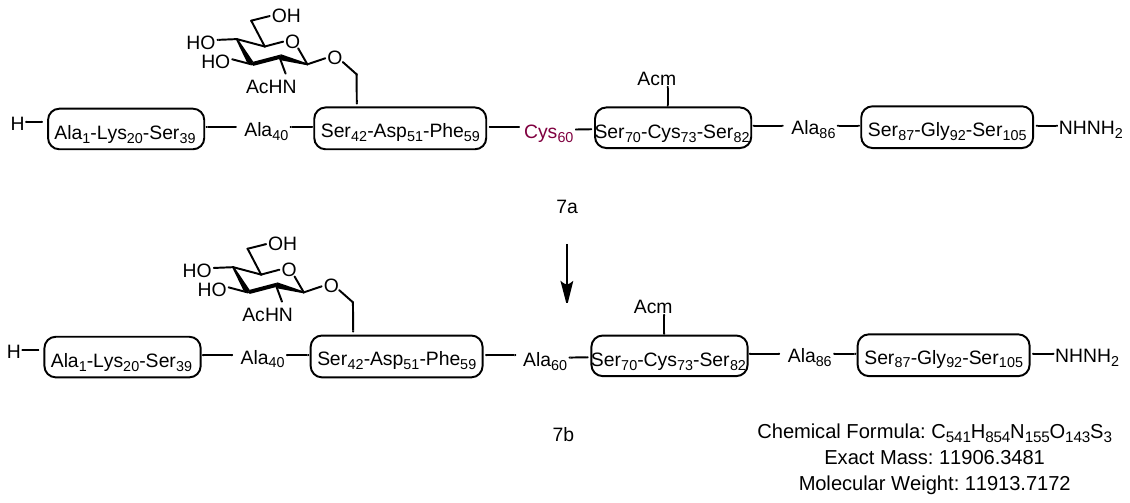

Peptide 7a (1.13 mg, 0.09 µmol) were subjected to the MFD conditions following General Procedure as described previously. The reaction was stirred at 37 ℃ for 5 hours, followed by addition of reducing buffer (6 M Gn·HCl, 200 mM Na_2_HPO_4_, 150 mM TCEP·HCl, pH 6.2) and quenched by CH_3_CN/H_2_O/AcOH (30/65/5, v/v/v). The product was purified using RP-HPLC (linear gradient 30-45% solvent B over 30 min, Proto-300 C4 column-1). Product was collected and lyophilized to provide peptide 7b (0.61 mg, 57%) as a fluffy white solid.

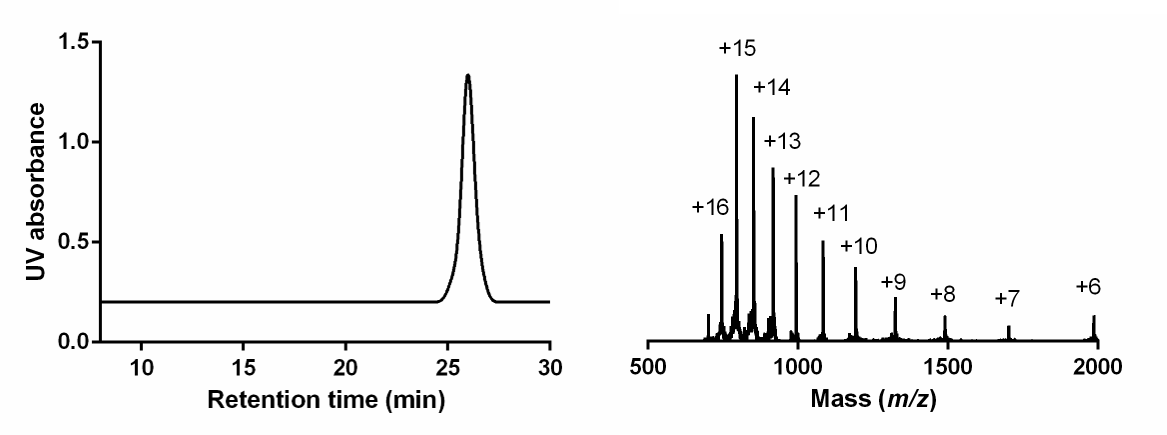

HPLC-MS analysis of **7b**. Left: UV traces (Liner gradient of 25-40% solvent B over 30 min, Proto-300 C4 column, tR = 19.6 min); Right: ESI-MS data. Calcd mass for C_541_H_854_N_155_O_143_S_3_: 11913.71 Da (average isotopes), [M+6H]^6+^ m/z =1986.61, [M+7H]^7+^ m/z =1702.95, [M+8H]^8+^ m/z =1490.21, [M+9H]^9+^ m/z =1324.74, [M+10H]^10+^ m/z =1192.37, [M+11H]^11+^ m/z =1084.06, [M+12H]^12+^ m/z =993.80, [M+13H]^13+^ m/z =917.43, [M+14H]^14+^ m/z =851.97, [M+15H]^15+^ m/z =795.24, [M+16H]^16+^ m/z = 745.60; observed: 1985.79, 1702.02, 1489.82, 1324.55, 1191.97, 1083.80, 993.63, 917.42, 851.74, 795.04, 745.62.

Peptidyl Hydrazide 9a

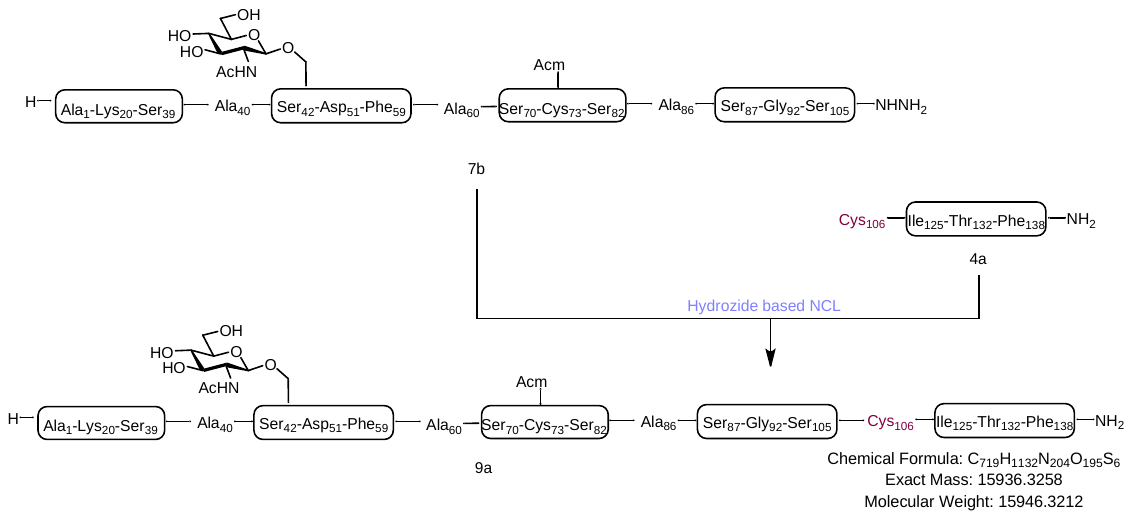

Peptidyl hydrazide 7b (2.31 mg, 0.20 µmol) and peptide 4a (1.14 mg, 0.28 µmol) were subjected to the ligation conditions following General Procedure as described previously. The reaction was stirred at room temperature for 12 hours, followed by addition of reducing buffer (6 M Gn·HCl, 200 mM Na_2_HPO_4_, 150 mM TCEP·HCl, pH 7.2) and quenched by CH_3_CN/H_2_O/AcOH (30/65/5, v/v/v). The ligated product was purified using RP-HPLC (linear gradient 25-45% solvent B over 30 min, Proto-300 C4 column-1). Product was collected and lyophilized to provide peptide 9a (1.02 mg, 34%) as a fluffy white solid.

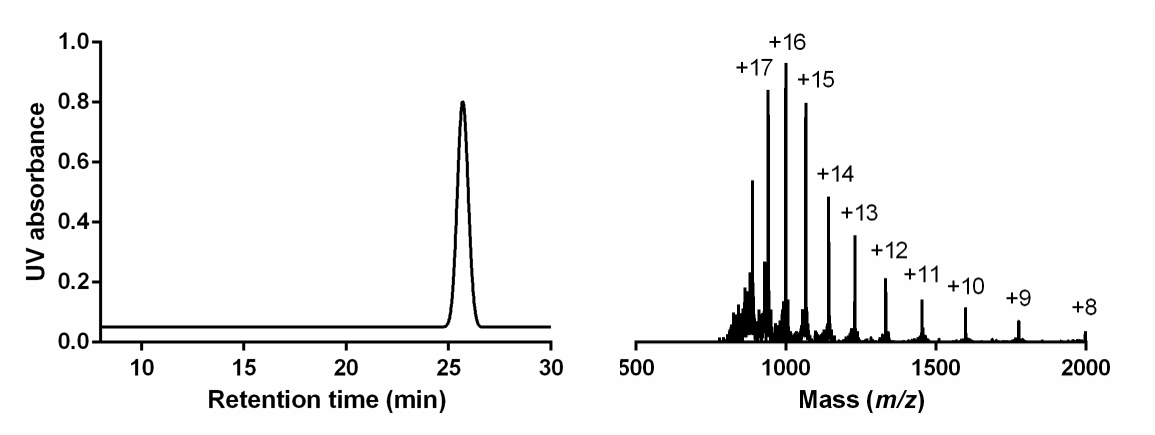

HPLC-MS analysis of **9a**. Left: UV traces (Liner gradient of 30-40% solvent B over 30 min, Proto-300 C4 column, t_R_ = 26.1 min); Right: ESI-MS data. Calcd mass for C_719_H_1132_N_204_O_195_S_6_: 15946.32 Da (average isotopes), [M+8H]^8+^ m/z =1994.29, [M+9H]^9+^ m/z =1772.81, [M+10H]^10+^ m/z =1595.63, [M+11H]^11+^ m/z =1450.66, [M+12H]^12+^ m/z =1329.86, [M+13H]^13+^ m/z =1227.64, [M+14H]^14+^ m/z =1140.02, [M+15H]^15+^ m/z =1064.08, [M+16H]^16+^ m/z =997.64, [M+17H]^17 +^ m/z =939.01; observed:1993.79, 1772.64, 1595.24, 1450.32, 1329.45, 1227.12, 1139.68, 1063.87, 997.15, 938.89.

Peptidyl Hydrazide 10a

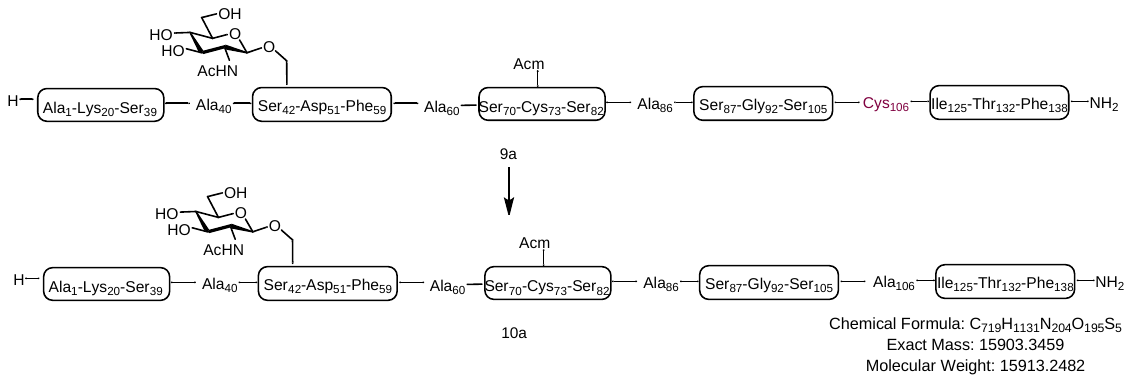

Peptide 9a (0.68 mg, 0.04 µmol) were subjected to the MFD conditions following General Procedure as described previously. The reaction was stirred at 37 ℃ for 5 hours, quenched by CH_3_CN/H_2_O/AcOH (30/65/5, v/v/v). The product was purified using RP-HPLC (linear gradient 25-45% solvent B over 30 min, Proto-300 C4 column-1). Product collected and lyophilized to provide peptide 10a (0.32 mg, 50%) as a fluffy white solid.

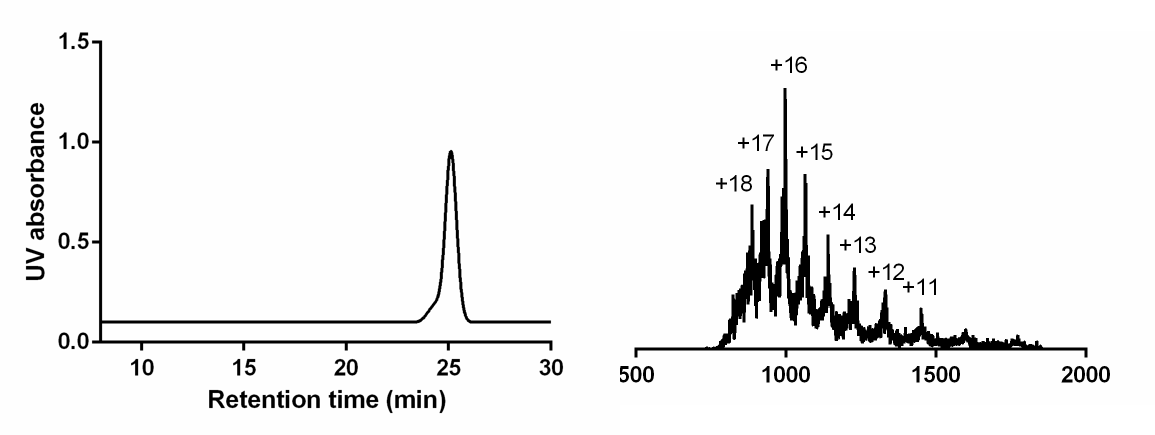

HPLC-MS analysis of **10a**. Left: UV traces (Liner gradient of 25-40% solvent B over 30 min, Proto-300 C4 column, t_R_ = 24.9 min); Right: ESI-MS data. Calcd mass for C_719_H_1131_N_204_O_195_S_5_: 15913.24 Da (average isotopes), [M+11H]^11+^ m/z =1447.65, [M+12H]^12+^ m/z =1327.10, [M+13H]^13+^ m/z =1225.09, [M+14H]^14+^ m/z =1137.66, [M+15H]^15+^ m/z =1061.88, [M+16H]^16+^ m/z =995.57, [M+17H]^17+^ m/z =937.07, [M+18H]^18+^ m/z =885.06; observed: 1447.56, 1326.71, 1225.05, 1137.42, 1061.04, 995.25, 937.20, 885.02.

Peptidyl Hydrazide 11a

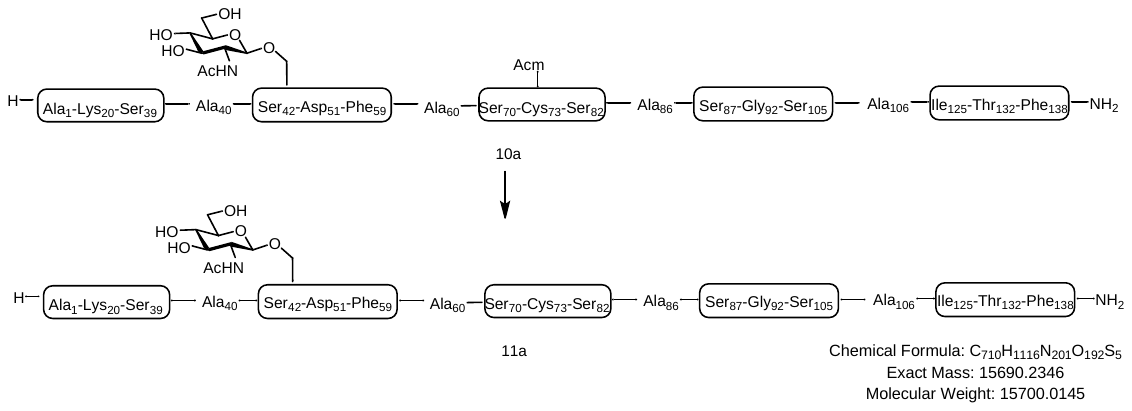

Peptide 10a (0.20 mg, 0.01 µmol) was subjected to the Acm removal conditions, quenched by CH_3_CN/H_2_O/AcOH (30/65/5, v/v/v). The product was purified using RP-HPLC (linear gradient 30-45% solvent B over 30 min, Proto-300 C4 column-1). Product collected and lyophilized to provide peptide 11a (0.11 mg, 61%) as a fluffy white solid

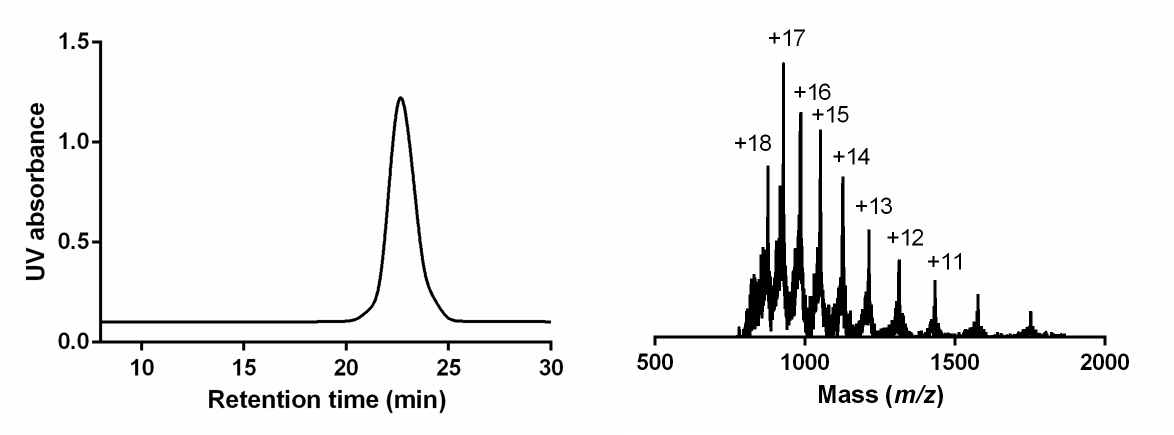

HPLC-MS analysis of **11a**. Left: UV traces (Liner gradient of 30-45% solvent B over 30 min, Proto-300 C4 column, t_R_ = 23.6 min); Right: ESI-MS data. Calcd mass for C_710_H_1116_N_201_O_192_S_5_: 15700.01 Da (average isotopes), [M+11H]^11+^ m/z =1428.27, [M+12H]^12+^ m/z =1309.33, [M+13H]^13+^ m/z =1208.69, [M+14H]^14+^ m/z =1122.42, [M+15H]^15+^ m/z =1047.66, [M+16H]^16+^ m/z =982.25, [M+17H]^17+^ m/z =924.53, [M+18H]^18+^ m/z =873.22; observed:1428.09, 1309.21, 1208.76, 1122.01, 1047.52, 982.06, 924.15, 873.31.

Peptidyl Hydrazide 12a

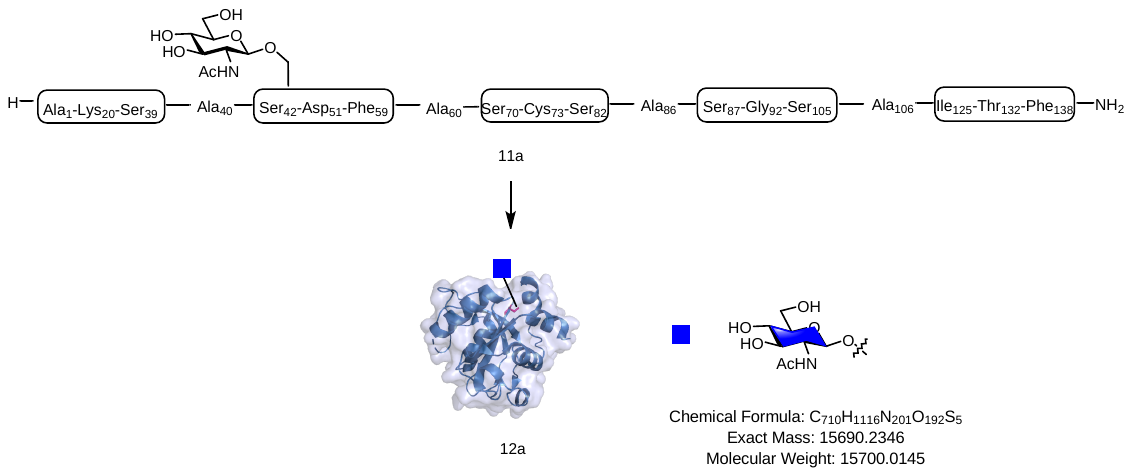

Protein 11a was dissolved in freshly prepared denaturing buffer (20 mM Tris-HCl, 500 mM NaCl, 10 mM DTT, 5% glycerol, 8 M urea, pH 7.5). The solution was then stirred at 4 ℃ with gradient dialysis against urea at decreasing concentrations to 0 M. The final solution was centrifuged through 3000 Da Amicon Ultra centrifugal filter devices (Millipore) at 4500 rpm to change buffer to Tris buffer (50 mM Tris-HCl, 100 mM NaCl, Ph 7.8), supernatant was collected and the concentration was determined by a NanoDrop 2000c (Thermo Fisher Scientific).

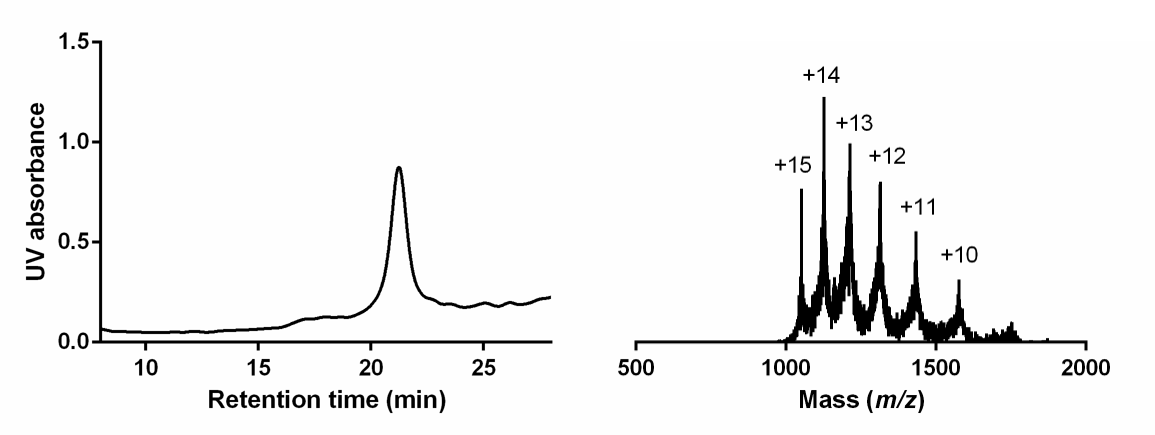

HPLC-MS analysis of **12a**. Left: UV traces (Liner gradient of 33-43% solvent B over 30 min, Proto-300 C4 column, t_R_ = 22.6 min); Right: ESI-MS data. Calcd mass for C_710_H_1116_N_201_O_192_S_5_: 15700.01 Da (average isotopes), [M+10H]^10+^ m/z =1571.01, [M+11H]^11+^ m/z =1428.27, [M+12H]^12+^ m/z =1309.33, [M+13H]^13+^ m/z =1208.69, [M+14H]^14+^ m/z =1122.42, [M+15H]^15+^ m/z =1047.66; observed: 1570.82, 1428.12, 1309.67, 1208.41, 1122.06, 1047.54.

Expression YTH domain **12b**

*E. coli* BL21 (DE3) cells transformed with the plasmids were plated on lysogeny broth (LB) agar plates containing specific antibiotics at 37 °C overnight. Single clones were selected and cultured in liquid LB medium containing specific antibiotics at 37 °C with shaking at 220 r.p.m. overnight. After 1:100 dilution in LB medium containing antibiotics, the culture was grown at 37 °C to an optical density at 600 nm (OD_600_) of 0.6–0.8. The culture was cooled to 16 °C and induced with isopropyl β-D-1-thiogalactopyranoside (IPTG) at 16 °C for 16–18 h. The cells were pelleted, resuspended in lysis buffer (50 mM Tris-HCl pH 7.8, 500 mM NaCl, 6 mM imidazole, 1 mM phenylmethylsulfonyl fluoride (PMSF)) and lysed. The lysates were centrifuged at 15,000*g* to remove the pellet. The supernatant was then purified with NTA-Beads and further purified by size-exclusion chromatography.

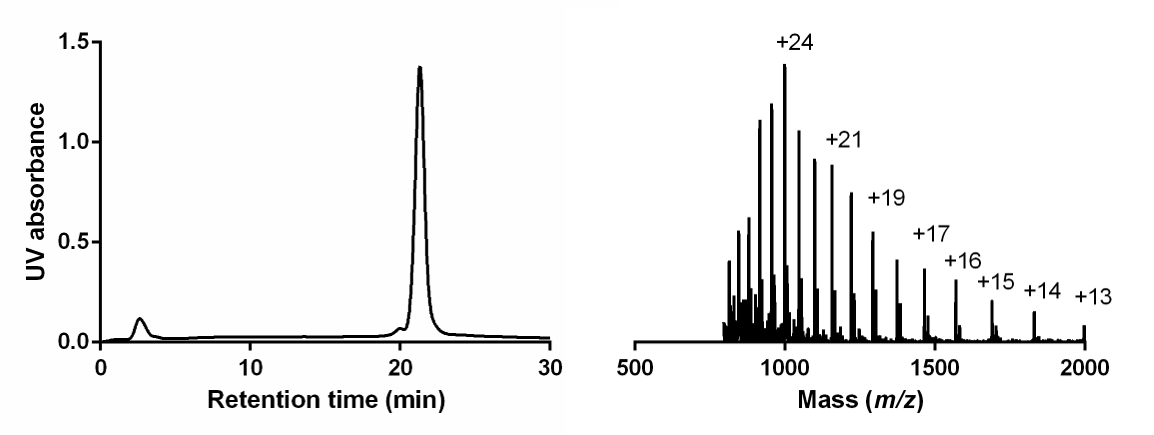

**Circular dichroism (CD) spectra of S396 YTH domain and S396 GlcNAcylated YTH domain**

Expressed YTH domain and synthesized S396 GlcNAcylated YTH domain were dissolved in water to a concentration of 0.2 mg/ml. The corresponding CD spectra were collected on a Jasco J-810 circular dichroism spectropolarimeter using a 1 mm pathlength cuvette at room temperature. The data were collected by running three scans from 190 nm to 250 nm at 0.2 nm intervals. Spectra were averaged and normalized against a baseline obtained by scanning water alone using the same parameters. The calculated results, based on the CONTINLL program from CDPro software, indicate that there is almost no difference for their secondary structure.

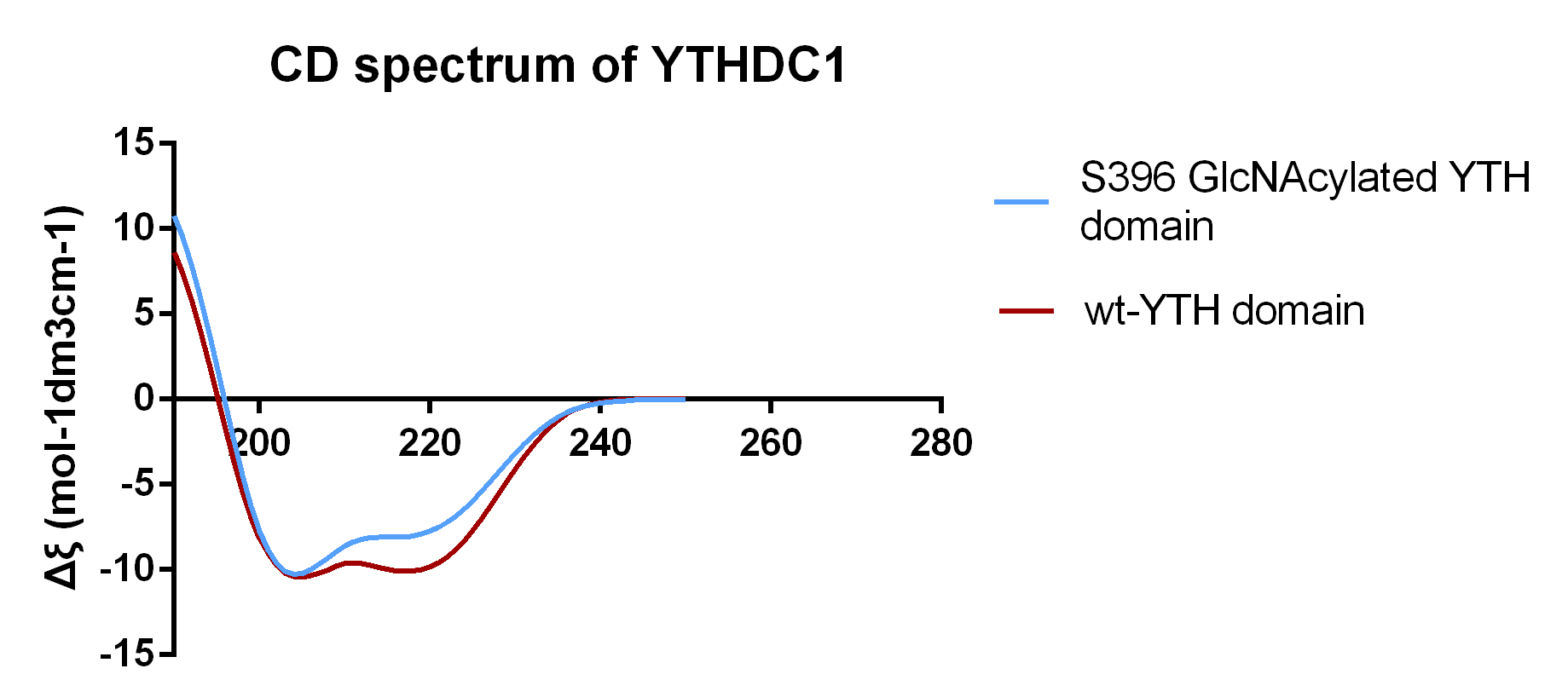

[1] Graham, M. E., Stone, R. S., Robinson, P. J. & Payne, R. J. Synthesis and protein binding studies of a peptide fragment of clathrin assembly protein AP180 bearing an O -linked β- N -acetylglucosaminyl-6- phosphate modification. Org. Biomol. Chem. 10, 2545–2551 (2012)

[2] Gude, M., Ryf, J. & White, P. D. An accurate method for the quantitation of Fmoc-derivatized solid phase supports. Lett. Pept. Sci. 9, 203–206 (2002)
